# Identification of SEC61B as a novel regulator of calcium flux and platelet hyperreactivity in diabetes mellitus

**DOI:** 10.1101/2024.02.20.581175

**Authors:** Yvonne X Kong, Rajan Rehan, Cesar L Moreno, Søren Madsen, Yunwei Zhang, Huiwen Zhao, Callum Houlahan, Siân P Cartland, Declan Robertshaw, Vincent Trang, Frederick Jun Liang Ong, Michelle Cielesh, Kristen C Cooke, Meg Potter, Jacqueline Stӧckli, Grant Morahan, Maggie Kalev-Zylinska, Matthew T Rondina, Sol Schulman, Jean Yang, G Gregory Neely, David James, Mary M Kavurma, Samantha Hocking, Stephen M Twigg, James Weaver, Mark Larance, Freda H Passam

**Author notes:** Correspondence Mark Larance, The Charles Perkins Centre, University of Sydney, John Hopkins Drive, Camperdown, Sydney, NSW 2050, Australia. Phone: Freda Passam, Haematology Research Lab, The Charles Perkins Centre, University of Sydney, John Hopkins Drive, Camperdown, NSW 2050, Sydney, Australia.

## Abstract

High platelet reactivity is associated with adverse clinical events and is more frequent in people with diabetes mellitus (DM). To better understand platelet dysfunction in DM, we performed a proteomic analysis in platelets from a matched cohort of 34 people without, and 42 people with type 2 DM. The cohorts were matched by clinical characteristics including age, sex, and coronary artery disease burden. Using high sensitivity unbiased proteomics, we consistently identified over 2,400 intracellular proteins, and detected proteins that are differentially released by platelets from people with diabetes in response to low dose thrombin. Importantly, we identified the endoplasmic reticulum (ER) protein SEC61 translocon subunit beta (SEC61B) was increased in platelets from humans and mice with *in vivo* hyperglycemia. SEC61B was increased in megakaryocytes in mouse models of diabetes, in association with megakaryocyte ER stress. A rise in cytosolic calcium is a key aspect in platelet activation, and the SEC61 translocon is known to act as a channel for ER calcium leak. We demonstrate that cultured cells overexpressing SEC61B have increased calcium flux and decreased protein synthesis. In accordance, hyperglycemic mouse platelets mobilized more calcium to the cytosol and had lower protein synthesis compared with normoglycemic platelets. Independently, *in vitro* induction of ER stress increased platelet SEC61B expression and markers of platelet activation. We propose a mechanism whereby ER stress-induced upregulation of platelet SEC61B leads to increased cytosolic calcium, potentially contributing to platelet hyperactivity in people with diabetes.

**Key Points:**

1. Platelet SEC61B is increased in hyperglycemia and contributes to increased endoplasmic reticulum (ER) calcium leak
2. Increased ER calcium leak is associated with ER stress and platelet hyperactivity

## Introduction

High platelet reactivity is the increased sensitivity of platelets to activate and secrete their content in response to various stimuli. Individuals with diabetes mellitus (DM) have high platelet reactivity both with and without treatment with antiplatelet agents.^1,2^ Clinically, this translates into a higher risk of cardiovascular events and impaired effectiveness to antiplatelet agents routinely used to treat coronary artery disease (CAD).^3,4^ Platelet activity largely relies on the secretion of stored intracellular proteins and metabolites, given their anucleate nature. Previous studies investigating the platelet proteome in people with diabetes have investigated only intracellular proteins in a limited number of patients.^5,6^ Unbiased analysis spanning the intracellular and secreted proteome can provide insights into the mechanisms contributing to high platelet reactivity.

In nucleated cells, newly synthesized membrane-bound and secreted proteins are translocated into the endoplasmic reticulum (ER) via the highly conserved SEC61 translocon, a heterotrimeric channel within the ER membrane.^7,8^ *De novo* protein synthesis has been described in platelets in response to *in vitro* activation and in pathological states, such as sepsis.^9,10^ Upon stimulation, platelets activate their splicing machinery to generate mature mRNAs, which can subsequently be translated into protein.^11^ Similar to nucleated cells, folding of membrane-bound and secreted platelet proteins is expected to take place in the ER, also known as the dense tubular system (DTS) in platelets. In addition, the platelet ER, or DTS, is an important calcium store and synthesis site of active lipids.^12^

Disruption of platelet ER homeostasis, by accumulation of misfolded proteins or depletion of calcium, causes ER stress which enhances platelet activation.^12^ ER stress is detected by intracellular sensors including inositol requiring protein-1 (IRE1) and protein kinase R-like ER kinase (PERK).^13^ Both IRE1 and PERK activation trigger signalling pathways of the unfolded protein response (UPR) aiming to restore ER homeostasis. Platelet ER stress was recently identified as a feature in patients with DM^14^, however, it is not known if this is linked to aberrant platelet protein synthesis or secretion.

In this study, we employed an unbiased proteomic approach to identify differences in platelet proteins in people with DM with suspected or known CAD compared with people with similar risk factors without DM. We provide the first comprehensive proteomic analysis of the intracellular and released platelet proteins in people with DM and CAD.

Our analysis identified that the beta subunit of the SEC61 transolocon (SEC61B) correlated with increasing serum fructosamine, a known measure of glycaemic control^15,16^. We show SEC61B upregulation in both platelets and megakaryocytes of hyperglycemic mice. Furthermore, increased expression of SEC61B in a cellular model and in platelets was associated with increased calcium leak from the ER into the cytoplasm, the development of ER stress, and a halt in protein synthesis. These results link SEC61B upregulation to ER stress and platelet activation, providing a mechanistic explanation of platelet hyperreactivity underlying CAD in patients with DM.

## Methods

*Details of human platelet aggregation, platelet proteomic studies, platelet and megakaryocyte immunofluorescence, in vitro studies of platelet ER stress, Western blot, flow cytometry, generation of SEC61B knockout and overexpression cell lines and calcium flux assays, are provided in the Supplementary Materials*.

### Patient cohort

A total of 76 people, 42 with type 2 diabetes mellitus (DM) and 34 without (non-DM), were recruited between 2020 and 2021 after informed consent, in accordance with the declaration of Helsinki. The study was approved by the Sydney Local Health District Human Ethics Committee (No X20-0085 & 2020/ETH00511).

The workflow of collection of clinical data and blood samples for the separation of plasma, serum and washed platelets is shown in **Fig 1A**. Clinical data including age, sex, body mass index, systolic and diastolic blood pressure, antiplatelet use, glycated haemoglobin (HbA1c), were derived from electronic medical records. Serum cholesterol, triglycerides, high density lipoprotein (HDL), low density lipoprotein (LDL) were analyzed in a routine chemical pathology lab. As HbA1c measurements were not available for all patients, serum fructosamine and glycated albumin were measured as markers of glycaemic control.^17^ Serum fructosamine was measured by spectrophotometry in a routine chemical pathology lab. Patient fructosamine was classified as high (> 290 µmol/L, approximately equivalent to HbA1c >7.0 %^15^) and normal (<290μmol/L, equivalent to HbA1c <7.0%) (normal range 200-290 µmol/L). Glycated albumin in plasma was measured by mass spectrometry for all patients.^18^ Coronary angiograms were reviewed and scored for Gensini and SYNTAX^19,20^ (as surrogate markers for coronary disease burden) by two interventional cardiologists and disagreement resolved by consensus (RR and JW).

**Figure 1.**
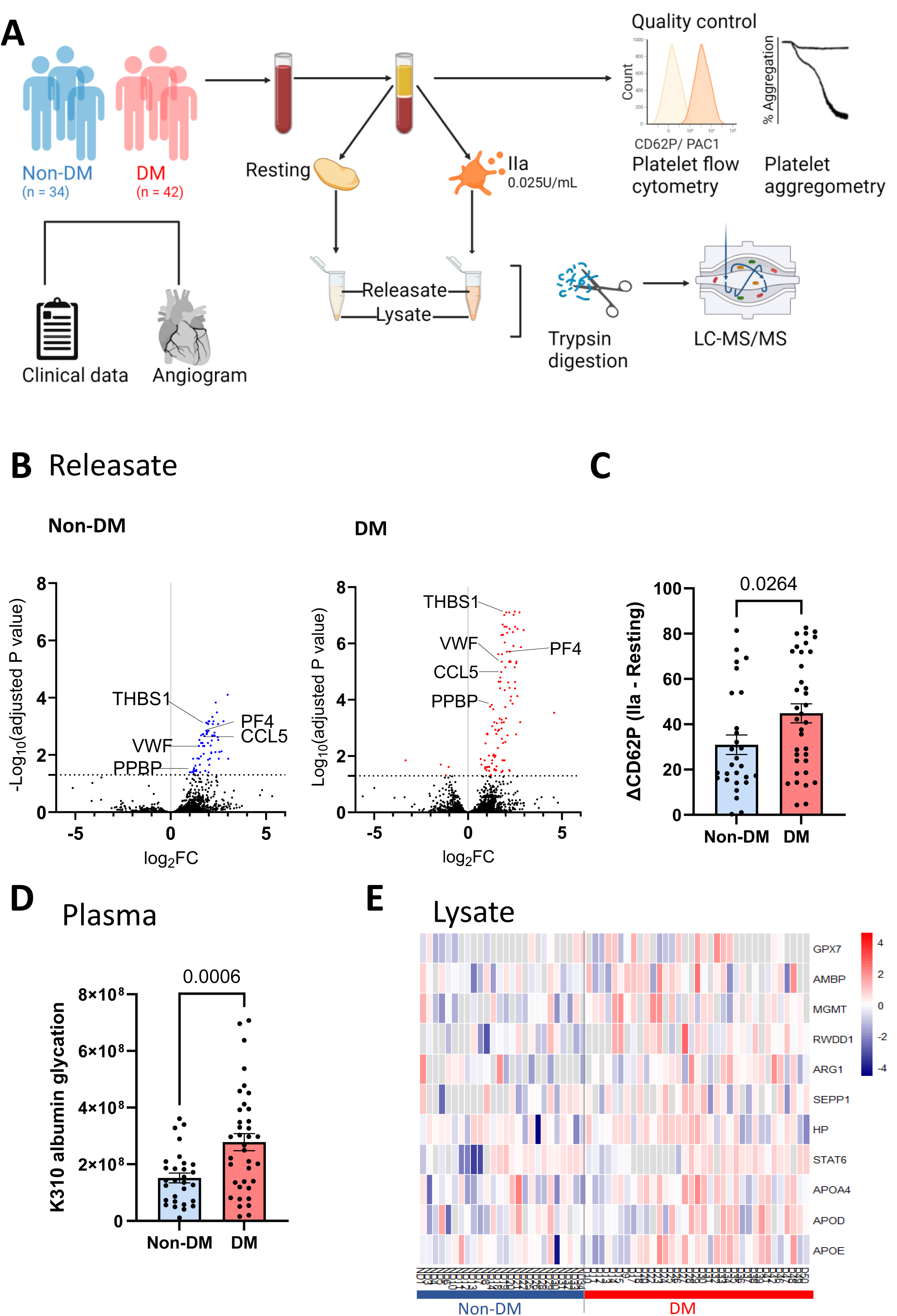
Proteomic analysis of platelet intracellular and released proteins from patients with or without diabetes and known or suspected coronary artery disease. **A.** Workflow demonstrating collection of clinico-laboratory and coronary angiogram data; quality check of platelets by flow cytometry; platelet aggregation; and, separation of resting and low-dose thrombin activated platelet intracellular fraction “lysate’ and released fraction “releasate”. **B.** Volcano plots of proteins detected in the releasate of non-diabetic (Non-DM) and diabetic (DM) platelets after stimulation with low dose thrombin 0.025 U/ml. Significantly released proteins from non-DM platelets are shown in blue and from DM platelets are shown in red. **C.** Difference of platelet surface CD62P expression before and after low dose thrombin (0.025U/mL) from patients with and without diabetes, Mann-Whitney test. **D.** Median intensity of K310 glycated albumin peptides detected by mass spectrometry in non-DM versus DM plasma, unpaired t-test with Welch’s correction. **E.** Heatmap of proteins involved in response to oxidative stress, detected in resting lysate proteins (Z-scores shown). 2467 proteins were consistently detected in >50% of samples. AMBP: alpha-1-microglobulin/bikunin precursor; APOA4: apolipoprotein A4; APOD: apolipoprotein D; APOE: apolipoprotein E; ARG1: arginase 1; CCL5:chemokine(C-C motif) ligand 5 (RANTES); GPX7: glutathione peroxidase 7; HP: haptoglobin; MGMT: O-6-methylguanine-DNA methyltransferase; PF4: platelet factor 4 (chemokine (C-X-C motif) ligand 4); PPBP: pro-platelet basic protein; RWDD1: RWD domain containing 1; SEPP1: selenoprotein P; STAT6: signal transducer and activator of transcription 6; THBS1: thrombospondin I; VWF: von Willebrand factor.

### Mouse models of diabetes

Eight- to nine-week-old male and female Apoe knockout (Apoe-/-) mice (Sydney Local Health District, Animal Welfare Committee, Australia, Protocol 2020-008) were treated with either citrate vehicle or streptozotocin (STZ, 75mg/kg) over five consecutive days via intraperitoneal injection (I.P.) (STZ: n = 3 male and n=2 female; Veh: n = 2 male and n=3 female). As female mice are resistant to the actions of STZ^21^, female mice received a second course 11 weeks after the first treatment course. Mice were fed a chow diet for 20 weeks before being sacrificed. Blood glucose was measured by glucometer (LifeSmart) prior to sacrifice.

Outbred mice were given ad libitum access to a high fat diet made in-house containing 45% fat, 35% carbohydrate and 20% protein from 10 weeks of age for 42 weeks. The characteristics and diet of these mice have been provided previously. ^22,23^ Experiments were performed in accordance with NHMRC (Australia) guidelines and under approval of the University of Sydney Animal Ethics Committee (protocol number #2017/1274 and #2021/1936).

Separately, for platelet immunofluoresence and calcium assays, male C57/BL6J mice were treated with STZ 55mg/kg I.P. daily for five consecutive days.^24^ They were fed with a chow diet and deemed hyperglycemic when the random blood glucose is confirmed to be greater than 15mM for at least two weeks prior to platelet isolation.

### Platelet calcium flux assay

Diluted mouse platelets were obtained by two rounds of serial centrifugation (240g, 2 minutes, soft brake) of mouse whole blood (1:15, whole blood: buffer) with HEPES-buffered Tyrode’s buffer (136.5mM NaCl, 2.68mM KCl, 20mM NaHCO3, 1.5mM Na2HPO4, 20mM HEPES, 5.55mM glucose) containing Cal-520-AM 2μM (Abcam, cat# ab171868) and probenecid 2mM (Sigma Aldrich). Diluted mouse platelets were incubated at 37°C in the dark for 30 minutes. The platelets were then diluted in an equal volume of HEPES-buffered Tyrode’s buffer and incubated at room temperature in the dark for 15 minutes to allow Cal-520-AM de-esterification. The diluted platelets were incubated with 1 mM calcium chloride for at least 3 minutes to allow the platelets to re-calcify. Subsequently, puromycin 100μM (Gibco, cat#A1113803), a protein synthesis inhibitor that maintains the SEC61 translocon in a calcium permeable state^25^, and EGTA 2mM were added, and a baseline was recorded for 60 seconds. Thapsigargin 2μM (Merk Life Science, cat# T9033-1MG) or vehicle (equal volume DMSO) were added, and events recorded for a further 240 seconds. A BD Accuri 6 flow cytometer was used with platelets recorded at the “slow” speed (14μl/min).

Human platelets were isolated from ACD-A anticoagulated whole blood (collected in BD Vacutainer ACD-A) as previously described.^26^ The platelet pellet was resuspended in HEPES-buffered Tyrodes buffer with BSA 0.35% (w/v) containing 2 µM fura-2-AM (Invitrogen), and incubated for 45 minutes in the dark. Subsequently, the fura-2-AM loaded platelets were pelleted by centrifuging at 700g for 5 minutes in the presence of platelet inhibitors apyrase 0.02U/mL (Sigma, cat # A6535-200UN) and prostaglandin E1 2µM (Cayman Chemical, cat# 13010-10mg). The pellet was resuspended in HEPES-buffered Tyrodes buffer with BSA and used immediately for calcium flux assays. The human platelets were incubated with 1 mM calcium chloride for 3 minutes immediately prior to the experiment. Then, the platelets were combined with 2 mM EGTA (and 1 µM puromycin in some experiments) and a baseline was recorded for 72 seconds prior to addition of sarcoendoplasmic reticulum calcium ATPase (SERCA) inhibitors (thapsigargin 2 µM; or BHQ 10 µM, 2,5,-Di-tert-butyl-1,4-benzoquinone, Sigma-Aldrich cat# 419648-5G) or vehicle control. The calcium flux was measured for a further 6 minutes after the addition of the SERCA inhibitors or control. Recordings for human platelet calcium flux experiments were performed using a BMG LabTech CLARIOstar plate reader.

### Platelet protein synthesis assay

For analysis of *de novo* platelet protein synthesis, 1µM L-azidohomoalanine was added to the HEPES-buffered Tyrode’s buffer and platelets were isolated from murine whole blood as above and incubated for eight hours. The platelets were allowed to adhere to a poly-D-lysine (Gibco, cat# A3890401) coated Nunc LabTek chamber slide. After the platelets were permeabilised, Click-iT AHA Alex Fluor 488 protein synthesis HCS assay (Invitrogen, cat# C10289) was performed as per manufacturer’s instructions. 15-20 platelets were analysed per animal to determine the mean L-AHA content per platelet per animal. The mean platelet L-AHA content was compared between normoglycemic and hyperglycemic animals using the Mann-Whitney test.

### Statistical analysis

Proteomic data were log2 transformed prior to further analysis. For the lysate proteome, label free quantification (LFQ) was used as it is anticipated the total protein content was unlikely to vary between patients. For the platelet releasate proteome, given minimal proteins are expected in the resting compared to thrombin-stimulated state, the assumptions required for LFQ analysis was not valid. Therefore, data were normalised against proteins that were minimally changed between the resting and thrombin stimulated states, which were typically plasma proteins (**Supplementary Table 1**). Preactivated platelet samples, in which the resting platelet CD62P expression was >20% on flow cytometry quality control, were excluded from the proteomics dataset in downstream analysis as we have previously described.^26^

Data were analysed using R (v 4.1.2) and visualised using the ggplot2 package and Graphpad Prism 10. Differential expression analyses between 1) samples from DM and non-DM, or 2) resting and thrombin-stimulated samples were conducted using moderated t-test in the limma R package.^27^ Gene Ontology analysis was conducted using the Gene Ontology Resource (http://geneontology.org/, accessed 03-10-2023).^28^ Wilcoxon rank test was used for analysis of samples pre- and post-treatment, such as for the ER stress markers after *in vitro* induction of ER stress in isolated platelets. Kruskal Wallis (with Dunn’s post hoc test) and Mann-Whitney tests were used for comparison of means in non-parametric samples. T-test with Welch’s correction for unequal variances were used for comparison of means in parametric data, after checking for normality using D’Agostino-Pearson omnibus, Anderson-Darling, Shapiro-Wilk, and Kolmogorov-Smirnov normality tests. Fisher’s exact test was used for comparison of proportions between multiple groups. Spearman correlation was used to correlate protein quantity and clinical characteristics, given the non-normal distribution of clinical parameters such as fructosamine, age, BMI, and coronary artery disease burden.^29^

## Results

To identify unique differences in the platelet proteome from patients with diabetes, people either with diabetes mellitus (DM) or without (non-DM) were recruited to the study and matched for age, sex, body mass index (BMI) and lipid profile. Patient characteristics are shown in **Table 1**. Coronary disease burden was quantified with two different angiographic scoring systems, the SYNTAX and Gensini scores. The Gensini score was 21.7 ± 22.4 in the DM group, and 11.2 ± 12.9 in the non-DM group. The SYNTAX score was 11.2 ± 12.9 in the DM group 6.7 ± 5.9 in the non-DM group. The differences in Gensini and SYNTAX scores were numerically higher in the DM group although not statistically significant. There were no differences in the proportion of patients on single or dual antiplatelet therapies between the DM and non-DM groups, nor was platelet aggregation in response to ADP or AA different despite antiplatelet use (**Table 1, Supplementary Fig 1A**).

**Table 1.** Characteristics of individuals included in the study. Values are given as average (± standard deviation). DM: diabetes mellitus; IIa: thrombin.

|  | **DM (n=42)** | **Non-DM (n=34)** | **P value** |
| --- | --- | --- | --- |
| Age (years) | 62.6 ± 10.5 | 66.0 ± 11.0 | 0.32 |
| Sex | M 80%; F 20% | M 66%; F 14% | 0.20 |
| Body Mass Index (kg/m<sup>2</sup>) | 30.0 ± 6.1 | 27.8 ± 4.8 | 0.31 |
| Platelet count (x10<sup>9</sup>/L) | 203 ± 42 | 213 ± 53 | 0.48 |
| Haemoglobin (g/L) | 143 ± 30 | 136 ± 23 | 0.84 |
| White cell count (x10<sup>9</sup>/L) | 6.9 ± 1.7 | 6.8 ± 1.8 | 0.80 |
| **Fructosamine** | **327 ± 114** | **255 ± 26** | **<0.0001** |
| SYNTAX score | 11.2 ± 12.9 | 6.7 ± 5.9 | 0.066 |
| Gensini score | 21.7 ± 22.4 | 16.7 ± 13.1 | 0.25 |
| ΔPAC-1 (%) after IIa stimulation | 27.1 ± 24.0 | 15.4 ± 18.6 | 0.18 |
| **ΔCD62p (%) after IIa stimulation** | **47.6 ± 24.3** | **34.4 ± 22.8** | **0.026** |
| Antiplatelet agents | None - 15<br>Aspirin - 14<br>P2Y12 inhibitor - 23<br>Dual - 13 | None - 9<br>Aspirin - 18<br>P2Y12 inhibitor - 18<br>Dual - 14 | 0.51 |
| Total Cholesterol (mmol/L) | 3.9 ± 0.8 | 4.0 ± 0.8 | 0.74 |
| Triglycerides (mmol/L) | 1.7 ± 1.0 | 1.4 ± 1.1 | 0.23 |
| HDL cholesterol (mmol/L) | 1.1 ± 0.3 | 1.1 ± 0.3 | 0.41 |
| LDL cholesterol (mmol/L) | 2.0 ± 0.8 | 2.2 ± 0.8 | 0.32 |
| Non-LDL Cholesterol (mmol/L) | 2.9 ± 0.9 | 2.9 ± 0.8 | 0.20 |

### Diabetic platelets have increased protein secretion in response to low dose-thrombin stimulation

To ensure a suitable comparison between DM and non-DM platelet responses, we used previously established titration studies on washed platelets from healthy donors. These determined that a low thrombin (IIa) dose of 0.025U/ml, results in submaximal platelet aggregation (40% by platelet aggregometry).^26^ We therefore treated washed platelets from all patients with 0.025U/ml thrombin to elicit potential platelet hyperreactivity to low dose thrombin. After thrombin stimulation a total of 109 and 71 proteins were significantly-secreted by DM and non-DM platelets, respectively (**Fig 1B**). Thrombin stimulation increased mobilisation of CD62P in platelets in the DM group when compared with the non-DM group (**Fig 1C)**, but the platelet aggregation between the two groups was similar (**Supplementary Fig 1A**).

Sixty-three proteins were released by platelets from DM and non-DM groups after thrombin stimulation (**Supplementary Table 2**). These included proteins involved in blood coagulation common pathway (GO:0072377), regulation of blood vessel remodelling (GO:0060312) and regulation of platelet activation (GO:0090330). Forty-six proteins were significantly released only by DM platelets. These included ADAMDEC1 (fold change 1.78 after IIa stimulation with p = 0.0015 for DM; compared with fold change 1.55 and p = 0.34 in non-DM), a soluble protease that cleaves platelet secreted pro-epidermal growth factor (pro-EGF) to high molecular weight EGF (HMW-EGF).^30^ HMW-EGF has been shown to contribute to increased thrombosis in an *in vivo* carotid injury model.^31^ Decorin (DCN; fold change 2.48 and p <0.0001 for DM and fold change 1.83 with p = 0.17 for non-DM), which has been shown to interact with integrin alpha2beta1 on platelets leading to platelet activation^32^, was also significantly secreted by DM but not non-DM platelets (**Supplementary Fig 1B**). As expected, analysis of patient plasma samples showed glycated albumin was higher in the DM group (**Fig 1D** and **Supplementary Fig 1C**).

### Enrichment of oxidative stress and ER stress pathways in DM platelets

Platelet lysate proteome analysis identified 2467 proteins across the samples. Proteins were ranked by fold change comparing DM and non-DM. The top 100 highest fold change differences between DM and non-DM platelet lysate proteomes in the resting state were subjected to Gene Ontology analysis. This revealed an enrichment for proteins involved in response to oxidative stress (**Fig 1E**), cellular oxidant detoxification and actin filament bundle assembly in line with oxidative stress being a feature of diabetes.^33^

### Positive correlation of platelet SEC61B abundance with serum fructosamine

From analysis of patient serum samples, fructosamine was significantly higher in patients with DM (**Table 1**). Correlation of the response to thrombin stimulation for the top 100 lysate proteins in DM compared to non-DM groups showed that only SEC61B demonstrated a significant positive correlation with serum fructosamine (**Fig 2A** and **Fig 2B**); but did not correlate with either SYNTAX (Spearman’s rho -0.011, p = 0.95) or Gensini scores (Spearman’s rho -0.064, p = 0.69). SEC61B is the beta subunit of the SEC61 translocon complex, which is important for polypeptide transport across the ER membrane.^34^ The SEC61 translocon is also a calcium “leak” channel from the ER.^35^ The quantity of SEC61B in the human platelet samples did not correlate with either of the other subunits of the SEC translocon (**Supplementary Fig 2A**) or with other ER marker proteins^36,37^ including CANX, CALR, GRP78 (HSPA5) and UGGT1, (p > 0.05 for all correlations with SEC61B) (**Supplementary Fig 2B**). Additionally, the quantity of GRP78 (HSPA5, also known as BiP) was similar between normal and high fructosamine platelets (**Fig 2C**), and non-DM and DM platelets (**Supplementary Fig 2B**), suggesting that the increase in SEC61B seen in hyperglycemia is independent of ER expansion.

**Figure 2.**
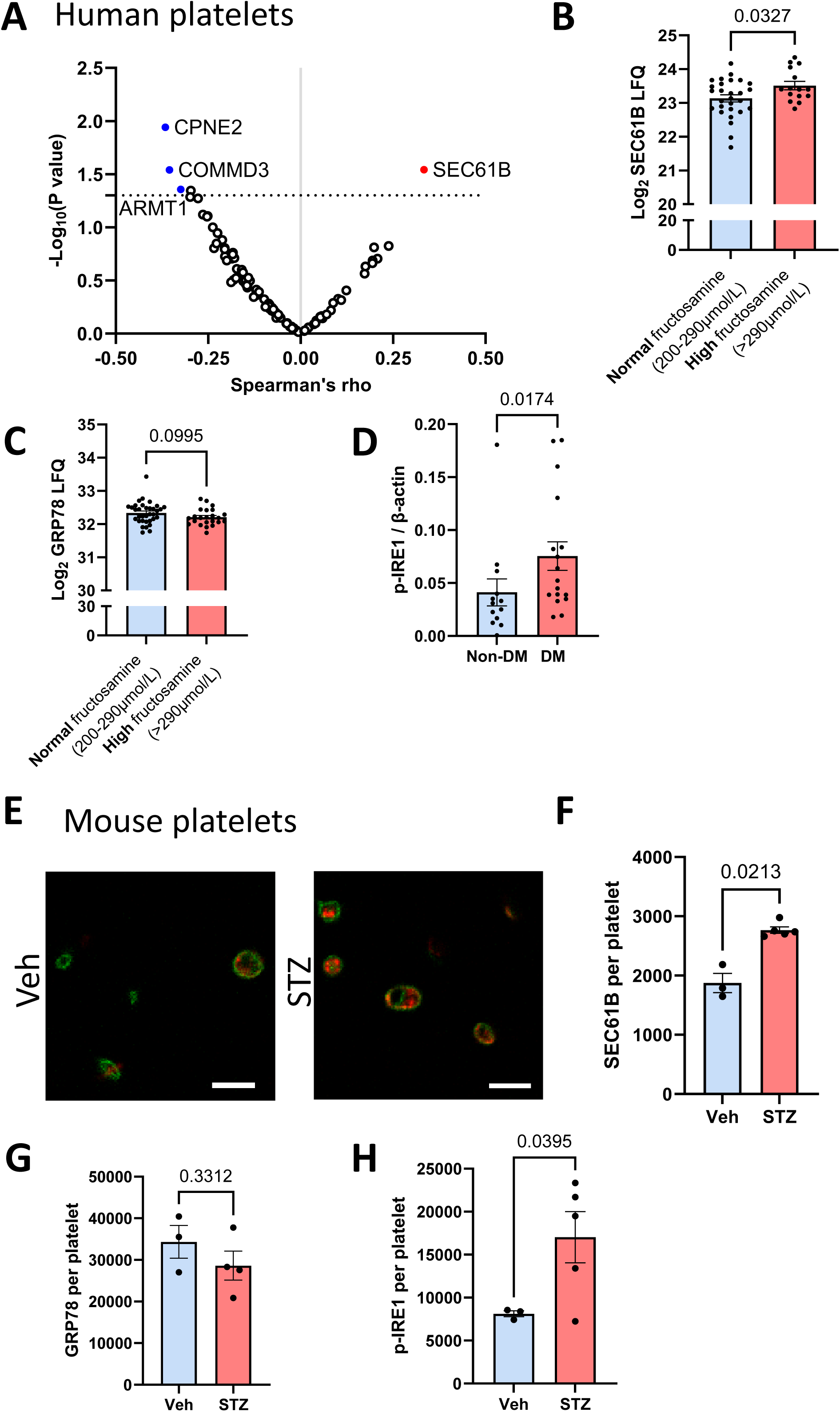
Increased platelet SEC61B protein is identified in hyperglycemic humans and mice. **A.** Correlation of the top 100 upregulated platelet lysate proteins, in response to thrombin, with serum fructosamine. SEC61B was the only platelet protein significantly correlated with serum fructosamine (red), Spearman’s rho = 0.33, p = 0.029. **B.** Log_2_ LFQ of SEC61B in platelets grouped in normal fructosamine (200-290 µmol/L) (blue) and high fructosamine (>290 µmol/L) (red), **C.** Log_2_ LFQ of GRP78 (also known as HSPA5 or BIP) in platelet lysates from patients without (non-DM, blue) or with diabetes (DM, red), t-test with Welch’s correction **D.** Ratio of phosphorylated IRE-1 (p-IRE1) to beta actin of platelet lysates from patients without (non-DM, blue) or with diabetes (DM, red) detected by Western blot. Mann-Whitney test. **E.** Platelets stained with antibody against SEC61B (red) and GP1Bb (green) isolated from normoglycemic (Veh) or hyperglycemic (STZ) C57BL/6 mice. Representative images. Scale bar 5 µm. **F.** Average SEC61B, **G.** GRP78 (HSPA5, BIP), and **H.** p-IRE1 fluorescence intensity of immunostained platelets from normoglycemic (Veh, n=3, blue) and hyperglycemic (STZ, n=4-5, red) mice. Average from n=15-20 platelets per mouse. Unpaired t-test with Welch’s correction. GP1Bb: glycoprotein 1B beta; GRP78: glucose regulated protein 78; HSPA5: heat shock protein A5, IRE1: inositol-requiring enzyme 1; Veh = vehicle, STZ = streptozotocin.

### Increased platelet SEC61B in hyperglycemia is associated with activation of IRE1 ER stress signalling pathway

The SEC61 translocon has previously been shown to attenuate ER stress signalling through the IRE1 pathway^38^, and recent evidence has identified platelet ER stress in DM.^14^ We therefore sought to investigate for evidence of platelet ER stress in our patient cohort. Phosphorylated IRE1 (p-IRE1), which is the active form of IRE1 generated upon ER stress, was significantly increased in DM platelets (**Fig 2D; Supplementary Fig 2C**), whereas platelet lysate IRE1 (official gene name ERN1) was comparable between non-DM and DM platelets (**Supplementary file “Platelet lysate proteins”**).

### Platelet SEC61B is increased in hyperglycemic mice

To determine if the correlation between hyperglycemia and increased SEC61B expression could be recapitulated in an animal model, we treated C57Bl/6J mice with streptozotocin (STZ) to induce hyperglycemia. The platelets for these mice were used for immunofluorescence studies and calcium flux assays. Hyperglycemic C57Bl/6J platelets had increased platelet SEC61B by immunofluorescence compared with control mice (**Fig 2E** and **Fig 2F**). Platelets from these mice had comparable levels of GRP78 (**Fig 2G**), but platelets from the hyperglycemic mice showed increased p-IRE1 at baseline (**Fig 2H**). These findings suggest that increased platelet SEC61B and p-IRE1 are independent of platelet ER expansion, as GRP78 is highly and ubiquitously expressed in the ER lumen.^39^

### Increase of SEC61B in hyperglycemia originates at the megakaryocyte level

To understand if upregulation of SEC61B in hyperglycemia originates from megakaryocytes, we employed different mouse models of DM for analysis of megakaryocyte SEC61B content. The first was injection of STZ in Apoe-/- mice^40^; and the second was an outbred mouse model fed on a high-fat diet^22^. The characteristics of the mice included in the study (Apoe-/- and outbred) are shown in **Table 2**.

**Table 2.** Characteristics of mice included in the study. Values are given as average (± standard deviation). BGL: blood glucose; STZ: streptozotocin; veh: vehicle.

| Mouse type | Average weight ± standard deviation (g) | Average fasting BGL ± standard deviation (mmol/L) |
| --- | --- | --- |
| Apoe<sup>-/-</sup><br>(n=5 vehicle; n=5 STZ) | STZ: 22.0 ± 1.0<br>Veh: 26.3 ± 2.8 | STZ: 11.9 ± 4.5<br>Veh: 5.6 ± 1.6 |
| Outbred<br>(n=5 normoglycemic; n=5 hyperglycemic) | Normoglycemic: 49.6 ± 14.7<br>Hyperglycemic: 63.9 ± 8.1 | Normoglycemic: 6.8 ± 1.2<br>Hyperglycemic: 16.0 ± 2.8 |

SEC61B was increased in the megakaryocytes of diabetic vs control Apoe-/- mice (**Fig 3A** and **Fig 3B**), and there was also an increase in the staining for p-IRE1 (**Fig 3C** and **Supplementary Fig 3A**). Similarly, immunostaining of bone marrow from the outbred mice showed increased SEC61B in hyperglycemic mice compared with normoglycemic mice (**Fig 3D** and **Fig 3E**) and an increase in the staining for p-IRE1 (**Fig 3F** and **Supplementary Fig 3B**). The increase in SEC61B appeared to be independent of ER expansion, as the ER chaperone GRP78 was unchanged in megakaryocytes of diabetic Apoe-/- (**Supplementary Fig 3C**) and hyperglycemic outbred mice (**Supplementary Fig 3D**), compared with their respective controls.

**Figure 3.**
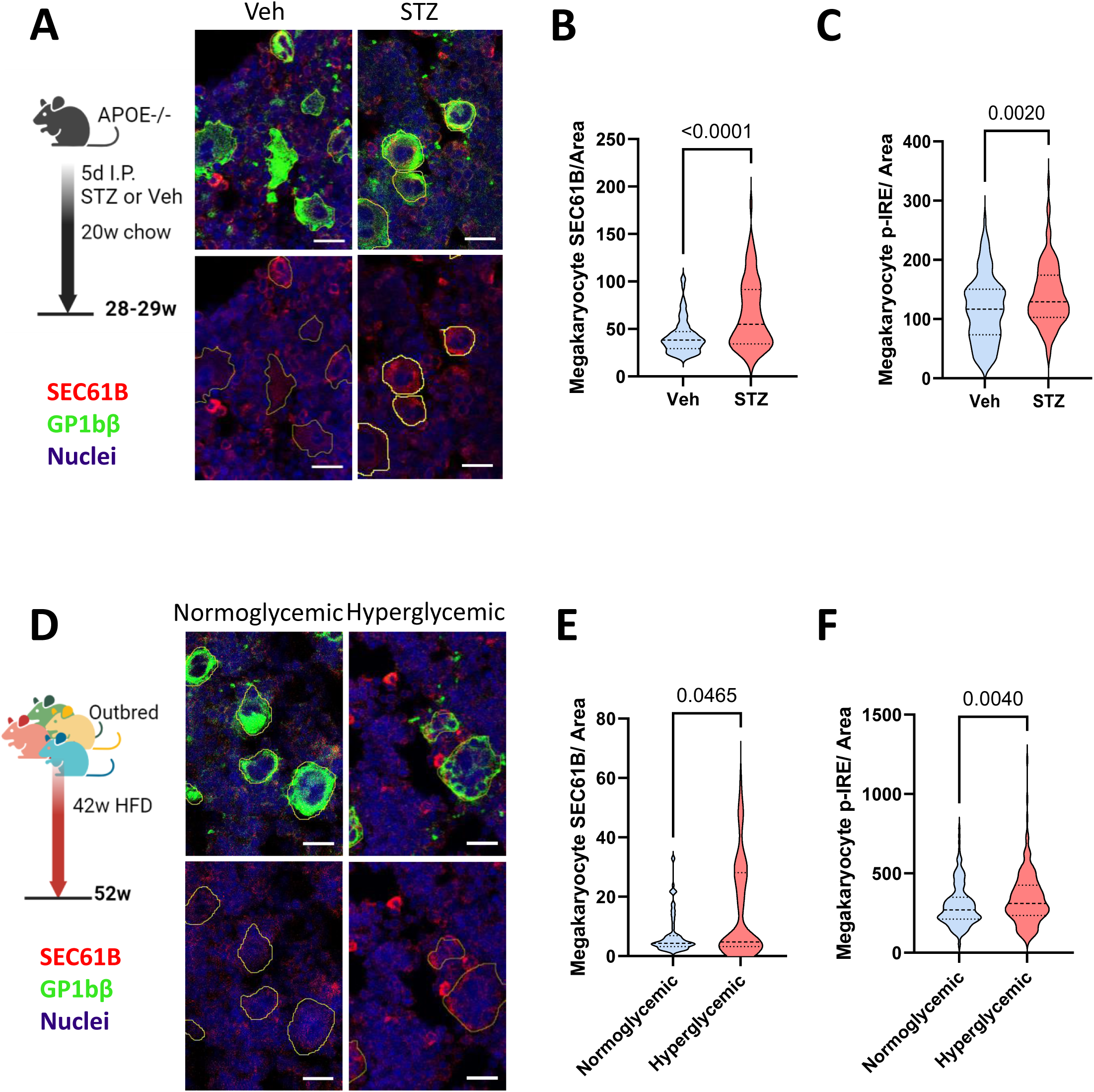
SEC61B is upregulated in megakaryocytes from murine models of diabetes mellitus. **A.** SEC61B fluorescence intensity in the bone marrow of Apoe-/- mice injected with Veh (left panel) or STZ (right panel), stained for SEC61B (red) and GP1bβ (green). Nuclei are stained with Hoescht 33258 (blue). Representative images. Scale bar 20µm. **B.** Mean megakaryocyte SEC61B fluorescence intensity per area and **C.** Mean megakaryocyte p-IRE1 fluorescence intensity per area of Veh or STZ treated Apoe-/- mice. **D.** SEC61B fluorescence intensity in the bone marrow of outbred mice with normoglycemia (left panel) or hyperglycemia (right panel) stained for SEC61B (red) and GP1bβ (green). Nuclei are stained with Hoescht 33258 (blue). Representative images. Scale bar 20µm. **E.** Mean megakaryocyte SEC61B fluorescence intensity per area and **F.** Mean megakaryocyte p-IRE1 fluorescence intensity per area of normoglycemic and hyperglycemic mice. n=5 mice for all groups. Mann-Whitney test.

### Overexpression of SEC61B in a cellular model is associated with increased calcium flux and decreased protein synthesis

Low basal cytoplasmic calcium is maintained by the function of sarcoendoplasmic reticulum calcium ATPase (SERCA) actively pumping calcium into the ER. Calcium “escapes” into the cytoplasm through leak channels, one of which is the SEC61 translocon.^35,41^ To interrogate the role of alterations in SEC61B levels in cellular calcium homeostasis, we employed genetic manipulation of SEC61B and chemical modulators of SEC61 and SERCA (**Fig 4A**). The drugs puromycin and eeyarestatin I (ES1) maintain the SEC61 channel in an “open”, calcium permeable state, to potentiate ER calcium leak.^25,42^ Puromycin inhibits polypeptide formation from the ribosome whereas ES1 binds to the lateral gate of SEC61 alpha subunit.^42,43^ Thapsigargin (TG) inhibits SERCA preventing the re-entry of calcium into the ER.^43^ The net effect of “opening” the SEC61 translocon and inhibiting SERCA is to induce maximal SEC61-mediated ER calcium leak (**Fig 4A**).

**Figure 4.**
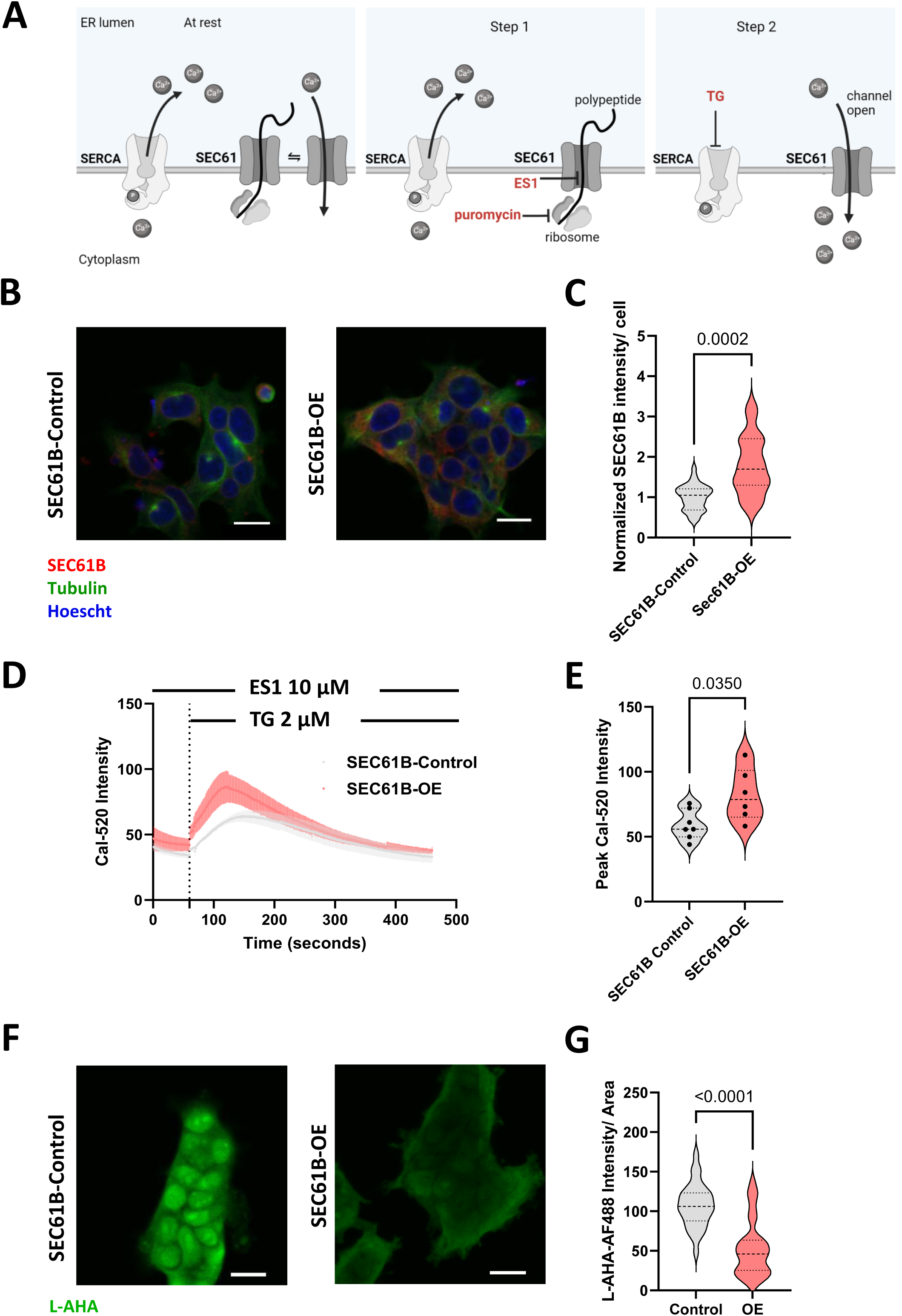
SEC61B overexpression in HEK 293 cells increased ER calcium leak and reduced cellular protein synthesis. **A.** Schematic for induction of SEC61 induced calcium leak with the combination of ES1 or puromycin followed by TG. **B.** SEC61B fluorescence intensity of lentiviral vector-transfected control or SEC61B overexpressing (OE) HEK293 cells. Cells were stained for SEC61B (red) and tubulin (green). Nuclei are stained with Hoescht 33258 (blue). Representative images. Scale bar 20µm **C.** SEC61B fluorescence intensity in control (grey) and OE HEK293 (red) cells (normalised to control), n=20-30 cells per. Unpaired t-test with Welch’s correction. **D.** Average fluorescence intensity of Cal-520 of control (grey) and OE (red) cells over time in response to ES1 and TG. (mean: solid line; SEM: shaded region), and **E.** peak ER calcium flux in control (grey) and OE cells (red) from n=5-6 independent experiments per genotype. Mann-Whitney test. **F.** L-AHA fluorescence intensity as a measure of protein synthesis, of lentiviral vector-transfected control or SEC61B OE HEK293 cells. Representative images. Scale bar 20µm **G.** L-AHA fluorescence intensity per cell area in control (grey) and OE (red) cells measured in n=30 cell clusters from n=3 independent experiments per genotype. Mann Whitney test. ES1: eeyarestatin I; TG: thapsigargin.

Transfection of HEK293 cells with a lentiviral SEC61B overexpression vector induced a 78.3 ± 72.4% increase in SEC61B expression compared with cells transfected with control vector (**Fig 4B, 4C**). Inducing ER calcium leak in HEK293 cells by treatment with ES1, followed by TG, resulted in significantly increased cytoplasmic calcium in SEC61B overexpressing cells compared with controls, 38.8 ± 34.1%, p=0.035 (**Fig 4D,4E**).

The SEC61 transolocon regulates protein synthesis by transporting peptides into the ER. *De novo* protein synthesis was determined by incorporation of L-azidohomoalanine (L-AHA). SEC61B overexpressing cells showed a significant decrease in protein synthesis by 51% compared with controls (**Fig 4F, 4G).**

ER calcium efflux was also significantly increased and *de novo* protein synthesis decreased in HEK293 cells with SEC61B depletion (**Suppl Fig 4A-E**). Thus, disrupting the stoichiometry of the SEC61 beta subunit appears to affect the function of the SEC61 translocon.

### Platelets from hyperglycemic mice have increased calcium flux and decreased protein synthesis

SEC61 translocon subunits have been reported in platelet transcriptomic^44^ and proteomic studies, but the function of the translocon in platelets has not been described. Using puromycin to maintain the SEC61 channel in a calcium permeable state, mouse platelets from hyperglycemic C57 BL/6J mice demonstrated greater ER calcium efflux with puromycin pre-treatment compared with normoglycemic C57 BL/6J mice, consistent with SEC61 translocon-mediated calcium leak (**Fig 5A-C**).

**Figure 5.**
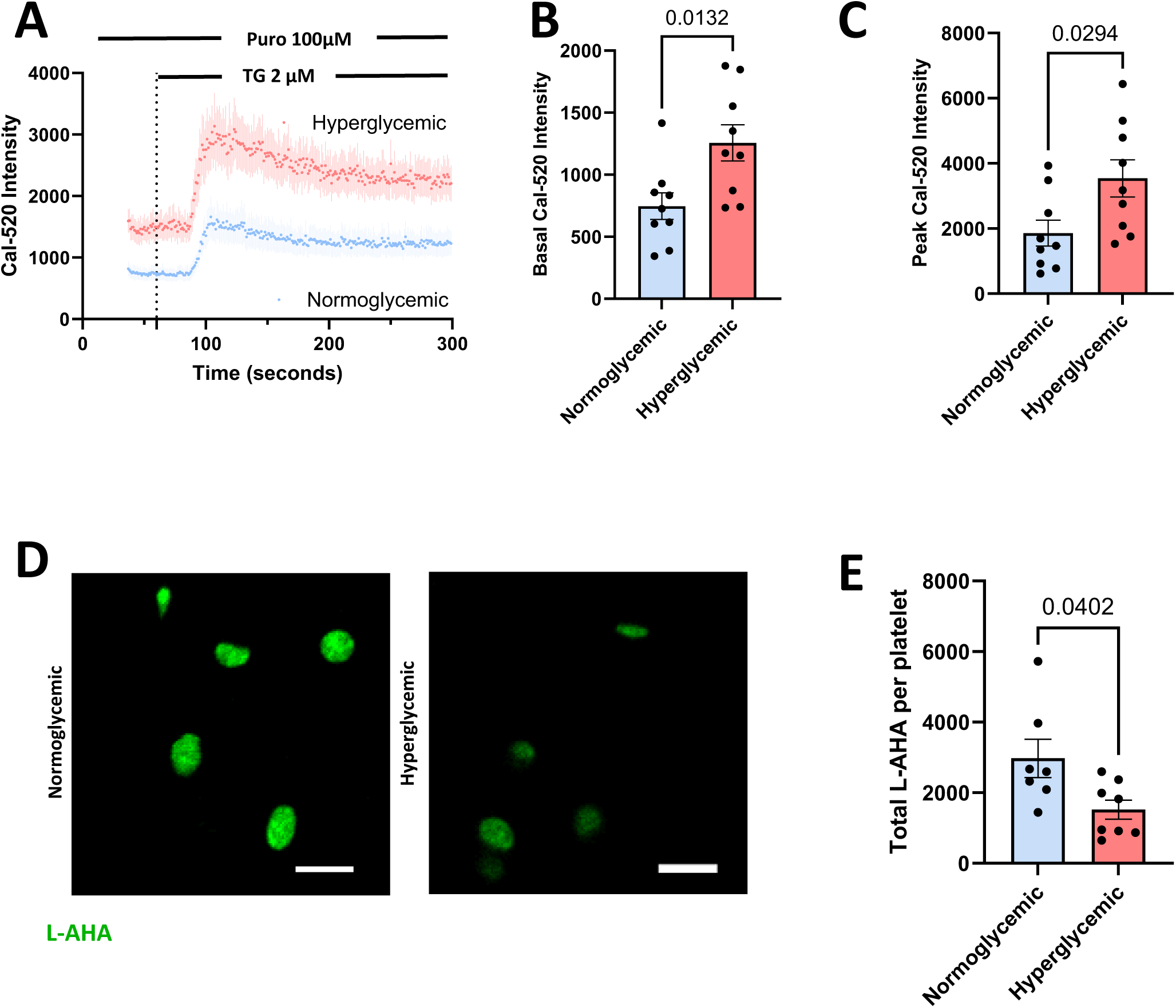
Diabetic mouse platelets have increased SEC61-mediated calcium leak and decreased protein synthesis. **A.** Cytosolic calcium over time measured by Cal-520 by flow cytometry in platelets from normoglycemic (n=8, blue) and hyperglycemic (n=9, red) C57 Bl/6J background mice. Platelets were treated with puromycin, followed by TG, to elicit SEC61-mediated ER calcium leak in the presence of EGTA. Bold line represents the mean and shaded line the SEM. **B.** Basal cytosolic calcium and **C.** peak calcium efflux from the ER in platelets from normoglycemic (blue) and hyperglycemic (red) mice. Unpaired t-test with Welch’s correction. **D.** L-AHA fluorescence intensity (green) of platelets from normoglycemic and hyperglycemic C57Bl/6J mice. Representative platelet immunofluorescence images. Scale bar 5 μm. **E.** L-AHA fluorescence intensity per platelet in normoglycemic (blue) and hyperglycemic (red) C57 Bl/6J background mice. Unpaired t-test with Welch’s correction. L-AHA: L-azidohomoalanine; Puro: puromycin; TG: thapsigargin.

We measured the incorporation of L-AHA in platelets from isolated from hyperglycemic C57 BL/6J mice. Platelets from hyperglycemic mice had decreased protein synthesis as measured by L-AHA incorporation *ex vivo,* compared with platelets from normoglycemic mice **(Fig 5D**, **5E**).

### Induction of ER stress in human platelets is associated with increased SEC61B and platelet secretion

To study SEC61 modulation by puromycin, we treated human platelets with puromycin followed by TG. This showed an increase in cytoplasmic calcium in response to puromycin and TG compared to TG alone (**Fig 6A**), suggestive of a role of SEC61-dependent permeability to ER calcium in platelets.

**Figure 6.**
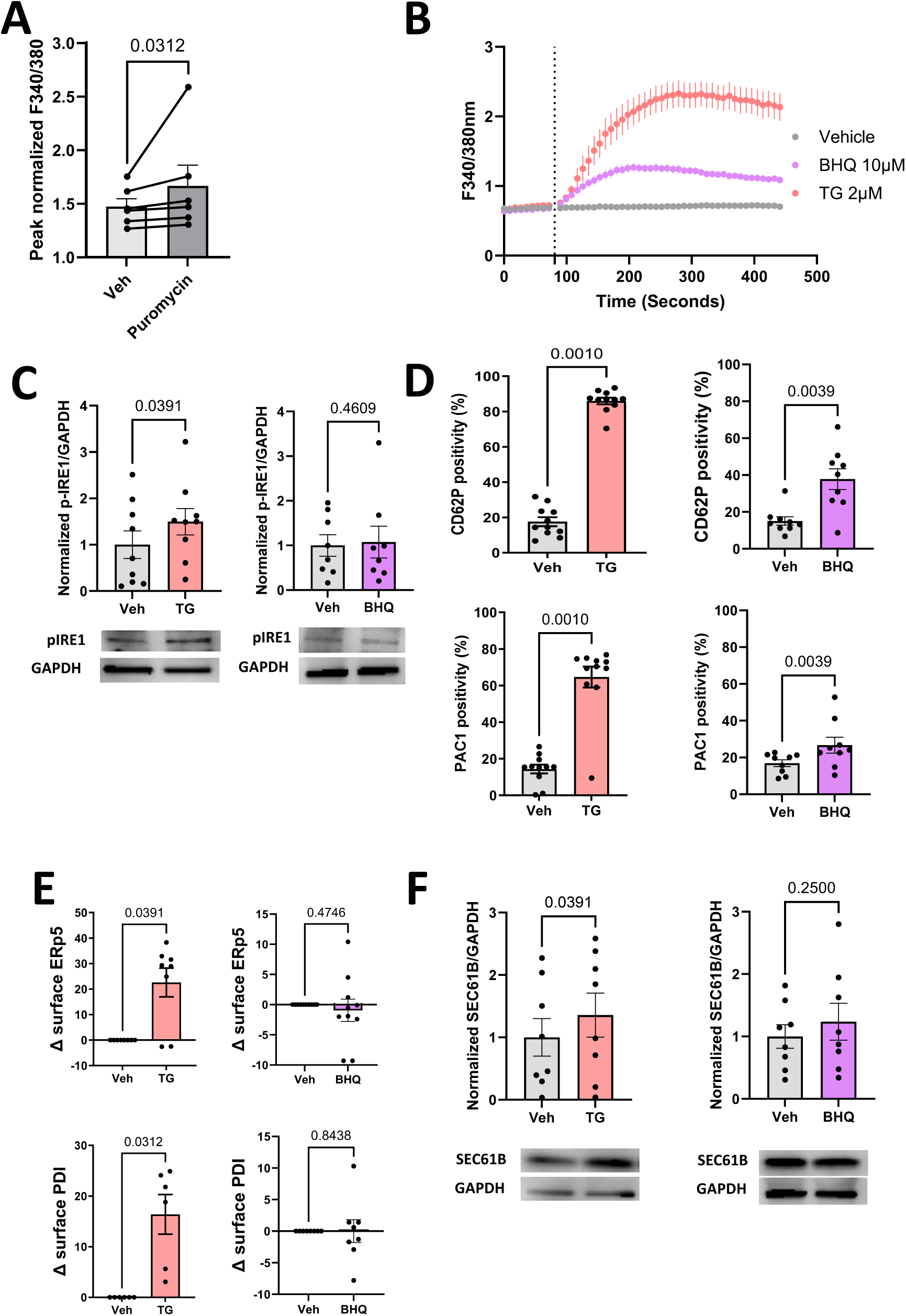
ER stress in human platelets increases platelet secretion and SEC61B expression. **A.** ER Calcium efflux is potentiated by puromycin in healthy human platelets. **B.** Time course of cytosolic calcium changes after TG (2μM, red) and BHQ (10µM, purple) in human platelets (n=3 healthy donors). **C.** Ratio of phosphorylated IRE-1 (p-IRE1) to GAPDH, **D.** CD62P and PAC-1, **E.** Surface PDI and ERp5, **F.** SEC61B to GAPDH band intensity of platelets treated with Veh (grey), TG (red), or BHQ (purple), Wilcoxon test. BHQ: 2,5,-di-t-butyl-1,4-benzohydroquinone; ERp5: endoplasmic reticulum protein 5; GAPDH: glyceraldehyde 3-phosphate dehydrogenase; IRE1: inositol-requiring enzyme 1; PDI: protein disulfide isomerase; TG: thapsigargin.

As we and others have shown that TG treatment induces ER stress in platelets^45^, we aimed to determine a threshold of cytoplasmic efflux for development of the ER stress response. For this we used two separate SERCA inhibitors: TG (inhibitor of SERCA2) and 2,5-di-(tert-butyl)-1,4-benzohydroquinone (BHQ) (inhibitor of SERCA3).^46^ Both TG and BHQ induced an increase in cytoplasmic calcium in human platelets (**Fig 6B**). However, only TG induced a platelet ER stress response through activation of the IRE1 pathway (**Fig 6C**). Neither TG nor BHQ affected the PERK pathway (phosphorylated eIF2a is downstream of PERK activation) (**Supplementary Fig 5A**). Both TG and BHQ stimulated the mobilization of alpha granule content as evidenced by increased platelet surface CD62P (**Fig 6D**). However, TG induced full activation of αIIbβ3 to a greater degree than BHQ as detected by the PAC-1 antibody (**Fig 6D**).

Furthermore, TG, but not BHQ, mobilized ER proteins, including protein disulfide isomerase (PDI) and endoplasmic reticulum protein 5 (ERp5) to the platelet surface (**Fig 6E),** but this was not associated with the concurrent release of the same proteins (**Supplementary Fig 5B**).

Stimulation of platelets with TG, but not BHQ, induced an increase in the expression of SEC61B (**Fig 6F**). The differential response to TG and BHQ supports platelet ER stress occurs with greater ER calcium loss and is associated with increased platelet secretion and activation (**Fig 5B-D**). Furthermore, the data support that SEC61B upregulation can occur *de novo* at the platelet level in response to platelet ER stress.

## Discussion

Increased platelet reactivity remains an urgent clinical issue in DM, as patients receive less benefit from traditional antiplatelet agents and are at increased risk of cardiovascular events. Here we provide the first comprehensive proteomic analysis of the intracellular and secreted platelet proteins in a matched cohort of patients with known risk of coronary artery disease, with and without DM. We identify increased platelet SEC61B in patients with recent hyperglycaemia. Functional studies suggest both increased and decreased SEC61B expression affect cellular calcium homeostasis, proteostasis, and platelet activation. We have thus identified for the first time that the SEC61 translocon plays a functional role in platelet biology and increased platelet SEC61B in DM can contribute to platelet hyperactivity.

Alterations in SEC61 alpha (SEC61A1) and gamma (SEC61G) subunits have been associated with disease.^47–50^ Altered SEC61A1 expression or function of SEC61A1 expression has been linked to diabetes mellitus, polycystic kidney disease, and immunodeficiency, whereas reduced SEC61A1 expression is associated with a decrease in ER calcium leak.^49,51,52^ Increased SEC61G has been reported in multiple solid organ malignancies, and is thought to promote cancer development, progression, and metastasis.^50,53–56^ However, descriptions of SEC61B leading to human disease^57^ are limited, and a role for the SEC61 translocon and the SEC61 beta subunit in platelet function are unknown.

We demonstrate SEC61B is upregulated in megakaryocytes and platelets from hyperglycaemic mouse models and platelets from humans with DM. We propose the increase in SEC61B is due to megakaryocyte and platelet ER stress. We demonstrated the *in vitro* induction of SEC61B in platelets by the ER stressor thapsigargin, and evidence of megakaryocyte ER stress in association with increased SEC61B. The SEC61B mRNA has been shown to closely associate with the ER, which may explain the rapid protein synthesis after platelet ER stress.^58^ XBP1s, the transcription factor downstream of IRE1 activation, has been shown to bind the SEC61B promoter in chromosomal immunoprecipitation assays.^59^ These observations support the potential for SEC61B induction both in the megakaryocytes, via transcriptional and translational upregulation, but also in the anucleate platelet via translational upregulation.

Overexpression of SEC61B in our cellular systems was associated with a decrease in protein synthesis. This was unsurprising given the important role of the SEC61 translocon in cellular proteostasis. Approximately one-third of all proteins are translocated into the ER for protein folding through the SEC61 translocon. ^34^ Whilst the SEC61A1 subunit forms the central pore for protein translocation, surrounding proteins including translocon-associated protein complex (TRAP complex) and GRP78 control and regulate this process.^41,60^ Therefore, whilst SEC61B and SEC61G are lateral to the SEC61A1 subunit, they may still play a regulatory role in protein synthesis. These peripherally placed subunits are thought to assist in translocon binding to the ribosome and other proteins involved in translation.^61,62^ Increased protein aggregates have been reported in diabetic platelets, consistent with platelet ER dysfunction.^14^ The reduced protein synthesis we observed in DM platelets may therefore be a response to ER stress, due to accumulation of protein aggregates, and altered properties of the SEC61 transolocon as a result of SEC61B upregulation.

The SEC61 translocon is permeable to calcium when the pore “breathes” after releasing a polypeptide.^34^ Decreased peptide transfer from decreased protein synthesis, in response to ER stress, may provide the opportunity for calcium to leak through the SEC61 channel into the cytoplasm. Although calcium movement through the translocon occurs via the SEC61A1 pore, SEC61B and SEC61G have been proposed to play a regulatory role.^61,62^ Previous studies have shown reduction in SEC61A1 is associated with decreased ER calcium leak.^51^ Although there is no information on the role of SEC61B in calcium flux, correct stoichiometry of channel subunits has been shown to affect the function of other channels. For example, the function of the ryanodine receptor (RyR), another ligand-activated calcium channel on the ER membrane, is regulated by other proteins that associate with the receptor. Depletion of calstabin-2 (FKBP12.6), a regulatory component of the RyR complex, and altered calstabin-2/RyR stoichiometry results in RyR instability and increased calcium leak from the receptor.^63^ Disruption of the SEC61 stoichiometry by upregulation of the beta subunit may thus underly increased permeability to calcium.

Platelets from people with DM have been described to have a higher basal cytosolic calcium compared to healthy age-matched controls.^64^ Oxidative stress has been shown to inactivate SERCA2 in platelets, with the reduced SERCA activity leading to elevated platelet cytosolic calcium.^65^ Our findings further these observations by implicating calcium leak via the SEC61 translocon as a possible additional mechanism of increased platelet cytosolic calcium in DM. This may provide a positive feedback cycle in which ER calcium depletion, as a result of SEC61B upregulation, further reinforces ER stress through activation of the IRE1 pathway. This then “primes” the platelet for subsequent activation (**Fig 7**).

**Figure 7.**
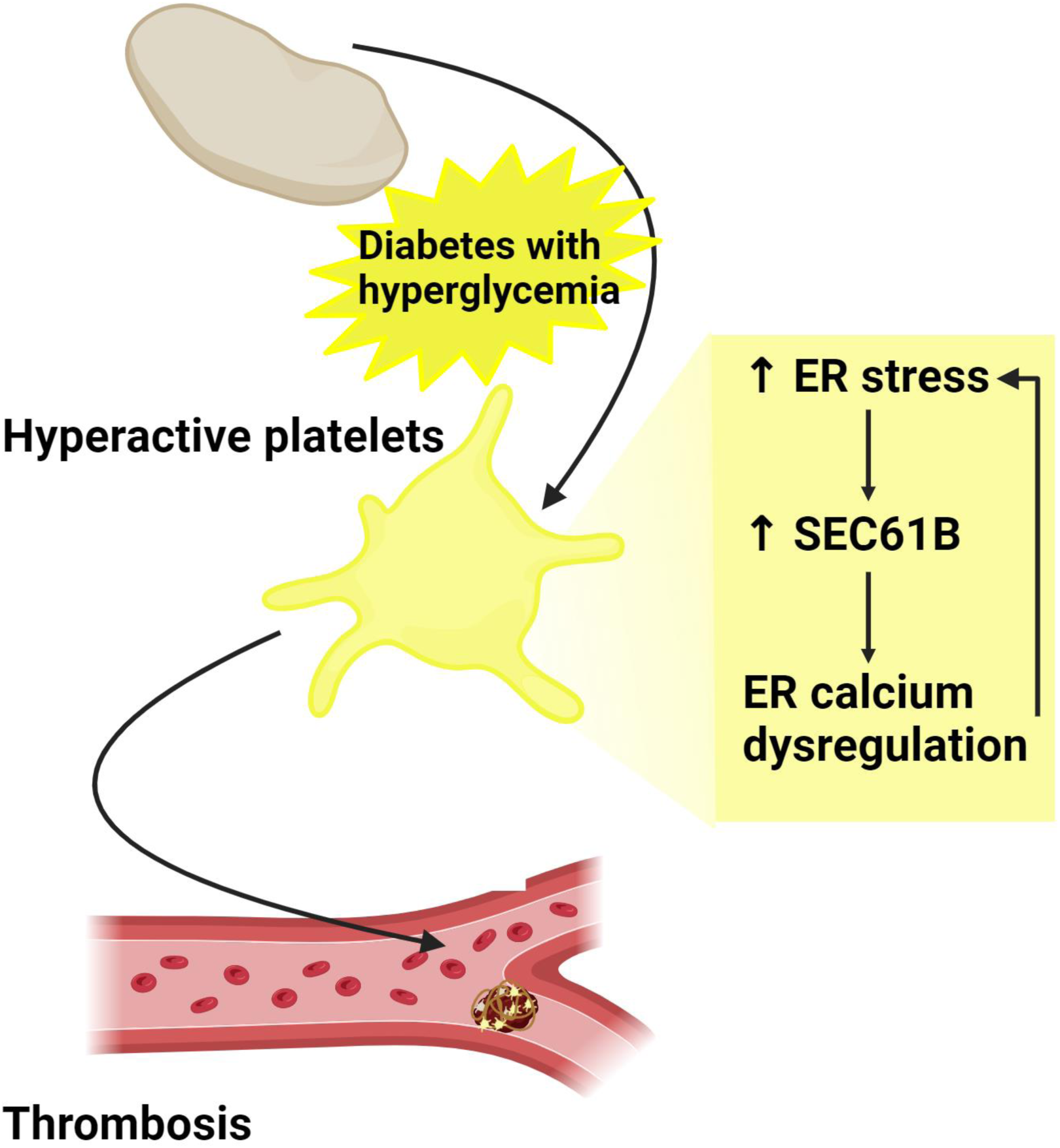
Proposed mechanism of platelet ER stress and increased SEC61B contributing to tendency for platelet activation in diabetes. Hyperglycemia in diabetes promotes ER stress with resultant SEC61B upregulation. Increased SEC61B provides a positive feedback cycle in which ER calcium depletion further reinforces ER stress. This then “primes” the platelet for subsequent activation.

In conclusion, we have identified SEC61B as a novel contributor to platelet hyperactivity in DM, which offers potential as a biomarker or for therapeutic targeting for the prevention of cardiovascular complications.

## Acknowledgements

We thank Prof Shaun P. Jackson, A/Prof Simone M. Schoenwaelder and Dr Yuping Yuan of the Thrombosis Group (Heart Research Institute); and Prof Peter Thorn (Charles Perkins Centre) for advice on calcium flux assays. We thank SydneyMS for providing the instrumentation used in this study.

Y.X.K. is supported by an NHMRC Postgraduate Scholarship and is a recipient of a Sydney Cardiovascular Initiative Catalyst Award in Precision Medicine 2020. M.L. is funded by a Cancer Institute New South Wales Future Research Leader Fellow. F.H.P. is recipient of a Sydney Cardiovascular Fellowship, University of Sydney and the Heart Research Institute, and a Ministry of Health NSW Cardiovascular Early Mid-Career Grant. This work was supported by a grant from MRFF Cardiovascular Health Mission Grant APP2017914 (to F.H.P and M.L) and a grant from the National Institutes of Health (HL142804 to M.T.R.).

## Authorship contributions

Y.X.K. designed and performed experiments, analyzed data, and co-wrote the manuscript. R.R. and J.W. recruited patients and contributed expertise. C.L.M, S.M., S.P.C. designed and performed experiments, contributed reagents and expertise. H.Z., C.H., D.R., V.T., F.J.L.O., M.C., M.P., performed experiments and analyzed data. K.C.C., J.S., G.M., G.G.N, D.J., M.M.K. contributed reagents and expertise. M.K.-Z., S.S., S.H., S.M.T., contributed to experimental design and provided expertise. Y.Z., J.Y. performed statistical analyses and provided expertise. M.T.R. contributed to experimental design and provided expertise. M.L. conceived the study, supervised research, designed experiments, provided expertise, and co-wrote the manuscript. F.H.P. conceived the study, supervised research, designed experiments and co-wrote the manuscript.

## Data availability statement

Proteomics datasets have been submitted to the PRIDE partner repository and accession number will be made available upon formal publication.

## Supplementary Materials

### Methods

Whole blood for platelet isolation was collected using a 19-gauge needle with light tourniquet into acid-citrate-dextrose (ACD-A) vacutainer tubes (Becton Dickinson, cat# 366645). Additional blood was collected into EDTA (Becton Dickinson, #367839) for full blood count indices and serum tube (Vacuette, cat#456010) for serum. Platelet isolation and quality control were performed as previously described.^26^

### Platelet aggregation

Platelet aggregation by light transmission aggregometry was performed on 300 uL platelet rich plasma for agonists ADP, arachidonic acid, collagen; and was performed on 300 uL washed platelets (400 x 10^3^/µL) in HEPES/Tyrodes buffer (HTGlc, consisting of 129 mM NaCl, 0.34 mM Na2HPO4, 2.9 mM KCl, 12 mM NaHCO3, 20 mM HEPES, 5 mM glucose, 1 mM MgCl2; pH 7.4) for thrombin (IIa) and U46619. Platelet aggregation was recorded for 10 min in response to ADP 5µM, arachidonic acid 0.5mg/mL, collagen 2µg/mL, IIa 0.025 U/ml, U46619 10 uM using an AggRAM 1484. Aggregation was determined as the peak light transmission expressed as % aggregation.

### Platelet releasate and lysate preparation for proteomic analysis

Double washed platelets (400×10^9^/mL in pre-warmed (37°C) HTGlc buffer, were divided into “resting control” and “thrombin stimulated” states. Apyrase 0.02U/mL and human thrombin 0.025U/mL (Sigma) were added to the resting control and thrombin treated samples respectively. All samples were incubated at 37°C for 5 minutes. PPACK 25nM (Abcam, cat# ab141451) was added to terminate thrombin action, and prostaglandin E1 (PGE1) 2µM was added immediately before centrifugation to obtain the platelet pellet. Platelet-released proteins (“releasate”) were in the supernatant, which was removed and stored under argon at -80°C. The platelet pellet was lysed to obtain the intracellular proteins (“lysate”) via resuspension in SDC lysis buffer (4% w/v sodium deoxycholate in 0.1M Tris-HCl pH 8.0) and stored under argon at -80°C.

### Platelet protein preparation for proteomic analysis

Concentration of proteins in the platelet releasate and lysate fractions were quantified using Bicinchoninic Acid (BCA) assay (Thermo Fisher). A total of 5 µg for platelet lysate, and 1 µg for platelet releasate, were processed as previously described for LC-MS/MS analysis.^26^ RAW proteomics data files were processed using the MaxQuant software (v 1.6.3.4) with reference to the MaxQuant contaminant and the human Uniprot (downloaded 5 May 2020) databases. A false discovery rate of 1% using a target-decoy based strategy was used for identification. The MaxLFQ algorithm was used for label-free quantification of proteins as previously described.^66^ Proteins detected in >50% of samples were included for downstream statistical analysis.

### Plasma proteomics analysis

Plasma samples were separated from the platelet poor plasma obtained after platelet isolation. Plasma (1 µL) was added to SDC buffer (24uL, 1% sodium deoxycholate, 10mM TCEP, 40mM chloroacetamide and 100mM Tris-HCl, pH 8.5) and heated for 10 minutes at 95°C. Subsequently, samples were left to cool to room temperature, diluted with water, followed by addition of LysC and trypsin (1:100 ratio for protease: protein, µg/ug). The samples were allowed to digest at 37°C for 16 hours and an equal volume of 99% ethyl acetate/1% trifluoroacetic acid (TFA) were added to the digested peptides. Digested peptides were loaded onto SDB-RPS StageTips to extract peptides for downstream analysis by LC-MS, as previously described.^18^ Proteins detected in >50% of samples were included for downstream statistical analysis.

### Platelet immunofluorescence

C57/BL6J or PDIA3-floxed on a C57/BL6J background were treated with citrate vehicle or streptozotocin (STZ, 55mg/kg) daily over five consecutive days via intraperitoneal injection. Mice were considered hyperglycemic with a random blood glucose level of > 15mM. 20uL of whole blood was collected via tail vein prick into 5uL of ACD. The whole blood was diluted in 150uL HEPES-buffered Tyrode’s buffer (20Mm HEPES, 134mM NaCl, 0.34mM Na_2_HPO_4_, 2.9mM KCl, 12mM HaHCO_3_, 1mM MgCl_2_, 5mM glucose, pH 7.4) with apyrase (0.04U/mL) and centrifuged at 270g in 37°C for 2 minutes. The supernatant containing the platelets were removed from the red and white cell pellet. This process was repeated by resuspending the pellet in Tyrode’s buffer. The supernatants were combined before being aliquoted into poly-D-lysine (0.1mg/mL) coated LabTek 8-well chamber slides. The platelets were allowed to adhere for at least one hour prior to fixation with 2% (v/v) paraformaldehyde. The platelets were washed with 3% BSA (w/v) prior to permeabilization with Triton-X100 (0.5%) in PBS for 20 minutes. Wells were blocked with 1% BSA (w/v) in PBS for at least one hour prior to incubation with SEC61B antibody (1:200 dilution, Cell Signaling Technology Cat #14648S); phospho-IRE1 (1:100 dilution, Thermo Fisher, cat# PA5105424); or GRP78 (1:200 dilution, Abcam, cat# ab21685) antibodies. Platelets were highlighted with anti-mouse GP1bβ -Dylight 488 (1:200, Emfret, cat # X488). Imaging was performed using a Zeiss 880 confocal microscope with an 100x oil objective. Approximately 15 to 20 platelets were analysed per animal using ImageJ analysis software, with the total target protein signal per platelet minus the signal in platelets without primary antibody but with the secondary antibody reported.

### *In vitro* induction of platelet endoplasmic reticulum (ER) stress and platelet flow cytometry

Washed platelets (400×10^6^/mL) were treated with thapsigargin (2µM) or BHQ (10 µM) for 1 hour or equal volume of DMSO control for *in vitro* induction of ER stress. Aliquots of the treated platelets were then analysed by flow cytometry for surface CD62P (BD Biosciences cat# 550888), PAC-1 positivity (Thermofisher Scientific Cat#MA528564), ERP5 (G-5 clone, Santa Cruz Biotechnology Cat #sc-365240 AF488) and PDI (Abcam Cat#ab202821-100uL). A BD Accuri 6 flow cytometer was used. The platelets were subsequently pelleted, lysed in RIPA lysis buffer with protease inhibitor cocktail (Sigma Aldrich, Cat#11873580001) and phosSTOP (Roche, cat#4906837001). The supernatant was resuspended in Laemmlli buffer with beta-mercaptoethanol and heated for 10min at 90°C for Western Blot analysis.

### Western blot

Platelet releasate or lysate samples were resolved on 4-20% polyacrylamide gels (Bio-Rad, cat# 0610374) under reducing conditions with beta-mercaptanol and transferred onto PVDF membranes using the IBlot2 Dry Blotting system (Thermo Fisher). PVDF membranes were blocked in 1% bovine serum albumin (BSA) in tris buffered saline-0.1% v/v Tween (TBS-T). The blots were incubated separately with primary antibodies overnight before secondary staining using a horse-radish peroxidase secondary antibody (1:1000). Primary antibodies used were anti-phosphoEIF2α (ser51) (1:1000, Abcam, cat # ab131505), anti-phospho-IRE1 (1:1000, Thermo Fisher, cat # PA5-105424), anti-PDI (1:100, clone DL-11, Sigma Aldrich), anti-SEC61B (1:1000, Cell Signaling Technology Cat #14648S), anti-GAPDH (1:4000, Thermo Fisher, cat#MA515738).

### Bone marrow immunofluorescence

Bilateral femurs from Apoe-/- and outbred mice were carefully dissected and fixed in 2% (v/v) paraformaldehyde for 24 hours. The femurs were decalcified in EDTA (0.5M) for 48 hours and dehydrated in sucrose (30% w/v) before being mounted in OCT and stored at -80°C. 10µm sections of bone marrow were obtained for immunofluorescence. Sections were blocked with foetal bovine serum (FBS, 10%), Triton-X100 (0.16%) in PBS for at least one hour prior to staining. Sections were stained with rabbit anti-phospho-IRE1 (1:100, Invitrogen, Cat#PA5-105424), rabbit anti-SEC61B (1:200, Cell Signaling Technology Cat #14648S), and rabbit anti-GRP78 (1:200 dilution, Abcam, cat# ab21685). Goat anti-rabbit IgG antibody conjugated to Alexa Fluor 647 (1:1000, Invitrogen, cat #A32733) was used to identify these markers. Megakaryocytes were identified by staining with anti-mouse GP1bβ - Dylight 488 (1:200, Emfret, Cat # X488) and nuclei were visualised by staining with Hoescht 33258 (1:10000). Imaging was performed using a Zeiss 880 confocal microscope with 40x water or 63x oil objectives. Images were taken to obtain ∼10-15 megakaryocytes per bone marrow sample. Image analysis was performed using ImageJ, with the mean intensity in the region of interest minus the mean intensity of megakaryocytes without primary antibody but with secondary antibody reported.

### SEC61B knockout and overexpressor HEK293 cell line generation

CRISPR knockouts were generating using the lentiCRISPRV2 vector (Addgene, cat# 52961). Lentiviruses carrying targeting sgRNAs or controls were packaged by co-transfection with pCAG-VSVG, and psPAX2 and the vector of interest at a ratio of 1:3:3 using lipofectamine 3000 in HEK293 cells. Virus was collected and purified from the media by centrifugation. For virus transduction, HEK293 cells were plated at 20% confluence, and exposed to the respective virus in the presence of 8ug/ml polybrene. Cells were then selected using puromycin (2μg/ml) for a period of 3 days. The sgRNAs sequences used for generating KO1 and KO2 were, 5’-ACCCCCAGTGGCACTAACGT-3’ and 5’-GTAGAATCGCCACATCCCCC-3’, respectively. The control sgRNA sequenced used was 5’-GCGTCTGAGATGAGAAAAGAT-3’.

The ORF for the Human SEC61B (NM_006808) was purchased from Origene (RG200247) and cloned using NEBuilder HiFi DNA Assembly (NEB #E2621) into a lentiviral backbone (Addgene, cat# 52961) for stable expression. The following primers were used to amplify the recipient vector 5’-atgaccgagtacaagcccac-3’ and 5’-GGTACCTTAATTAACCAAACTGGATC-3’. Two fragments were amplified and inserted for rational vector design. Fragment 1, SEC61B ORF, was amplified using these primers 5’-ATCCAGTTTGGTTAATTAAGGTACCTGAATCAATATTGGCAATTAGCC -3’ and 5’-GGACAGTGCCAAGCAAGCA-3’; Fragment 2, Ef1a promoter, was amplified from Addgene 52961, using 5’-TTGAGTTGCTTGCTTGGCACTGTCCttttttgaattcgctagctaggtcttgaaag-3’ and 5’-cgcaccgtgggcttgtactcggtCATatggtggcagcgctctagaac-3’. Control viruses without SEC61B ORF insert were also generated. Viruses were packaged in HEK293 cells with psPAX2 (Addgene #12260), pCAG-VSVg (Addgene #35616) using Lipofectamine 3000 Transfection Reagent (Thermo Fisher Scientific). HEK293 cell lines were infected in the presence of polybrene (8μg/ml) and selected for 1 week in 2μg/ml of puromycin before proceeding to experiments.

Quantification of SEC61B expression in the control, knockout and overexpressor cells were determined by measuring the SEC61B quantity per cell in ImageJ, with an average of 30 cells measured per genotype.

### HEK293 calcium flux assays

HEK293 cells, with SEC61B overexpression or control, were plated onto fibronectin-coated (25 μg/mL) Nunc LabTek 8-well chamber slides at 75,000 cells per well and allowed to recover overnight. The cells were incubated with Cal-520-AM (Abcam, cat# ab171868) 2μM, probenecid (Sigma-Aldrich) 2mM and Pluronic F127 (Sigma-Aldrich) 0.1% v/v final concentration, in phenol-red free DMEM supplemented with 10% fetal calf serum (FCS) for 30 minutes at 37°C, 5% CO_2_. The Cal-520-AM loading solution was removed and replaced with fresh phenol-red free media supplemented with 10% FCS, and cells were further incubated for 15 minutes at 37°C, 5% CO_2_ to allow the Cal-520-AM to de-esterify. The media was then removed, and cells washed with calcium free imaging buffer (125 mM NaCl, 2 mM MgCl_2_, 4.5 mM KCl, 10 mM glucose, 20 mM HEPES, pH to 7.4). Eeyarestatin I (ES1, Cayman Chemical cat# 41260-54-4) 10 μM was added immediately prior to the calcium flux assay to permit the translocon-mediated calcium leak from the ER (1x volume). Calcium mobilisation from the ER to the cytosol was induced addition of 3x volume of thapsigargin solution (thapsigargin, Merk Life Science cat# T9033-1MG, dissolved in imaging buffer, final thapsigargin concentration 2 µM) after 75 seconds of baseline readings. Cells were imaged with a Zeiss 880 confocal microscope with 20x lens using the “Time Course” function with images recorded every second. The data were analysed with ImageJ using the “Time Series Analyzer” plugin. A total of 4-6 regions of interest were identified per experiment, and these were used to generate the mean Cal-520 signal per experiment. The mean Cal-520 signal for each experiment were used to generate the time course and for the peak Cal-520 signal analysis.

HEK293 cells, with SEC61B knockout or control, were seeded into poly-D-lysine (Thermo Fisher, cat#A3890401) coated 96 well clear bottom black plates at 75,000 cells per well. The cells were allowed to recover for ∼24 to 48 hours prior to the calcium flux assays. HEK293 cells were incubated with fura-2-AM (Thermo Fisher, cat# F1201) 2µM with pluronic F127 (0.1% v/v final concentration) and probenecid 2mM in phenol red free DMEM (Gibco) supplemented with 10% FCS and GlutaMAX 1x (Gibco) for 60 minutes at 37oC, 5% CO2. The fura-2-AM loading solution was removed and replaced with fresh phenol red free media supplemented with 10% fetal calf serum (FCS) and cells were further incubated for 30 minutes to allow for fura-2-AM de-esterification. Eeyarestatin I (ES1) 10uM, which captures SEC61 complex in Ca^2+^-permeable state^42^, was added immediately prior to the calcium flux assay to potentiate translocon-mediated calcium leak from the ER. Calcium mobilization from the ER to the cytosol was induced by automated injection of thapsigargin (final concentration 2µM), mixing by automated double-orbital function after 90-120 seconds of baseline readings. The fluorescent readings were conducted using the BMG Labtech CLARIOstar Plus with the cytosolic calcium represented by the ratio of fluorescent emission at 510 nm, after excitation at 340 nm and 380 nm. ‘

### HEK293 protein synthesis assay

HEK293 cells with SEC61B knockout or overexpression, and their respective controls, were seeded onto poly-D-lysine coated Nunc Lab-Tek II chamber slides at 75,000 cells per well. The cells were allowed to recover for ∼24 hours prior to the protein synthesis assay. The Click-iT AHA Alex Fluor 488 protein synthesis HCS assay (Invitrogen, cat# C10289) was used according to the manufacturer’s instructions, with the alteration that cells were incubated with media containing L-azidohomoalanine (L-AHA) for 1 hour. Cells were imaged using a Zeiss 880 confocal microscope with 100x oil objective. Images were taken to obtain ∼30 cellular clusters per genotype. Image analysis was performed using ImageJ, with the mean intensity in the region of interest minus the mean-intensity of cells treated with media without L-AHA reported.

## Supplementary Results

### Platelet aggregation responses show variability in patients with or without diabetes

Platelet aggregation results are shown in **Supplementary Fig 1A**. Response to ADP was supressed in both non-DM and D groups reflecting use of P2Y12 antagonists in both groups. Despite the use of aspirin there remained patients with > 50% aggregation in response to arachidonic acid indicative of decreased response to aspirin. Similarly, there were patients with hyperresponsive platelets to low dose thrombin and U46619. Response to collagen was blunted in both groups.

### SEC61B knockout in HEK 293 cells increased ER calcium leak and reduced cellular protein synthesis

After puromycin clone selection, SEC61B expression was reduced by 41.5-77.7% in the SEC61B knockout (KO) cell lines (p ≤0.003, **Supplementary Fig 4A**). ER calcium efflux was significantly increased with SEC61B depletion (mean 8.6±5.4% increase for KO-1 and 12.0±9.0% increase for KO-2 compared with KO-control, p = 0.0068; **Supplementary Fig 4B** and **4C**).

The KO cells had a lower rate of *de novo* protein synthesis as determined compared to the control cells (mean 57.7±20.4% reduction for KO-1 and 71.9±16.9% reduction for KO-2 compared with KO-control, p<0.0001; **Supplementary Fig 4D** and **4E**).

---

**Supplementary Table 1.** Proteins used as controls for normalisation of platelet releasate proteins.

|  |  |  |  |  |  |  |
| --- | --- | --- | --- | --- | --- | --- |
| A1BG | APOC3 | CA1 | HPR | IGJ | KRT9 | SERPINA6 |
| A2M | APOD | CD9 | HPX | IGKC | KRT10 | SERPINC1 |
| ACTC1 | APOE | CD36 | IGFALS | IGKV2-40 | KRT14 | SERPIND1 |
| AFM | APOL1 | CP | IGHA1 | IGKV3D-11 | KRT16 | SERPING1 |
| AGT | ARF1 | DCD | IGHG1 | IGKV3D-20 | ORM1 | SLC4A1 |
| ALB | ARPC2 | FGA | IGHG2 | IGKV4-1 | ORM2 | SPTLC2 |
| AMBP | AZGP1 | FGB | IGHG3 | IGLL5;IGLC1 | PARVB | STOM |
| APOA1 | BIN2 | FGG | IGHG4 | ITGB3 | PON1 | SYNE1 |
| APOA2 | C4A | GC | IGHM | KRT1 | PPIB | TF |
| APOA4 | C4BPA | GP1BA | IGHV3-15 | KRT2 | RAP1B | TUBB |
| APOC1 | C5 | HBA1 | IGHV3-66 | KRT5 | RSU1 |  |
| APOC2 | C9 | HP | IGHV3-72 | KRT6A | SERPINA3 |  |

**Supplementary Table 2.** Proteins released by DM and non-DM platelets after low dose (0.025U/mL) thrombin stimulation.

| Non-DM |  |  |  | DM |  |  |  |  |  |
| --- | --- | --- | --- | --- | --- | --- | --- | --- | --- |
| ANG | ECM1 | MAN2A1 | SRGN | ADAMDEC1 * | CTSC * | FUT8 * | MMRN1 | QSOX1 | TIMP3 * |
| ANGPT1 | EFEMP1 | MMRN1 | ST3GAL6 ** | ANG | CTSD | GALNT1 * | NID1 | RARRES2 | TNXB * |
| APLP2 | EMILIN1 | NID1 | ST6GAL1 ** | ANGPT1 | CTSW | GALNT2 * | NID2 | RNASE4 * | TPMT * |
| APP | ESAM | NID2 | TFPI | APLP2 | CXCL3 | GGH * | NIT1 * | RNASET2 * | TPP1 |
| B4GALT1 ** | F5 | NUCB1 | TGFB1 | APP | CXCL5 | GLCE * | NUCB1 | SDC4 * | TREML1 * |
| C8B | FAM3C | PCSK6 | THBS1 | AQR * | DAG1 | GLG1 | NUCB2 * | SEC23IP | UXS1 * |
| CALU | FGA | PDGFA | TIMP1 ** | BDNF * | DCN * | GOLIM4 * | PCOLCE * | SERPINE1 | VEGFC |
| CCL5 | FGB | PDGFB | TPP1 | C8B * | DNAJC3 | GP5 | PCSK6 | SERPINE2 | VWF |
| CLEC11A | FGG | PDGFD | VEGFC | C8G * | DPP7 | HPSE | PDGFA | SIAE * | XYLT2 |
| CPB2 ** | FIS1 ** | PF4 | VWF | CALU | ECM1 | IGF2 | PDGFB | SORT1 * |  |
| CST3 | FSTL1 | PPBP | XYLT2 | CCL5 | EFEMP1 | IGFBP2 * | PDGFD | SPARC |  |
| CTGF | GLG1 | PROS1 |  | CFHR2 * | EMILIN1 | ISLR * | PEPD * | SPP2 |  |
| CTSA | GP5 | PSAP |  | CLEC11A | ESAM | LAMC1 * | PF4 | SPTA1 * |  |
| CTSD | HEXB ** | QSOX1 |  | CLSTN1 * | F5 | LOXL3 | PLG * | SRGN |  |
| CTSW | HPSE | RARRES2 |  | CLU * | FAM3C | LTBP1 | PLOD3 * | STAT6 * |  |
| CXCL3 | IGF2 | SEC23IP |  | COCH * | FGA | LYZ * | PLXDC2 * | TFPI |  |
| CXCL5 | LGALS3BP ** | SERPINE1 |  | CSDE1 * | FGB | MAN1A1 | PPBP | TGFB1 |  |
| DAG1 | LOXL3 | SERPINE2 |  | CST3 | FGG | MAN1B1 * | PRKDC * | TGFBI * |  |
| DNAJC3 | LTBP1 | SPARC |  | CTGF | FN1 * | MAN2A1 | PROS1 | THBS1 |  |
| DPP7 | MAN1A1 | SPP2 |  | CTSA | FSTL1 | MAP3K5 * | PSAP | TIMP1 |  |

\* Indicates proteins that are uniquely released by DM platelets after low dose (0.025U/mL) thrombin stimulation; \*\* indicates proteins uniquely released by non-DM platelets after low dose thrombin stimulation.

## Supplementary Material

### Figure Legends

**Supplementary Fig 1.**
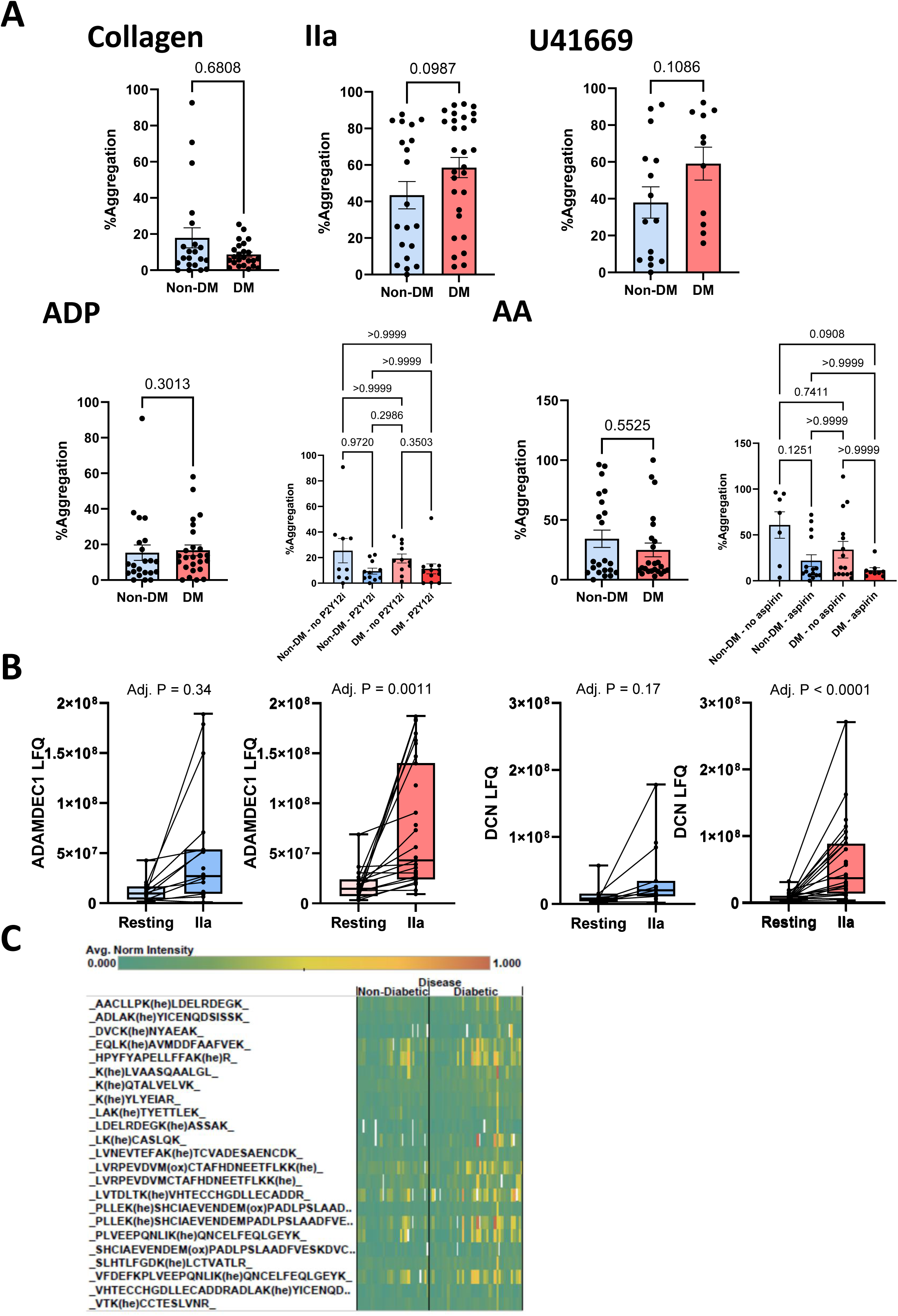
Platelets from patients with diabetes show increased secretion of prothrombotic proteins despite similar aggregation patterns compared with patients without diabetes. **A.** Platelet aggregation expressed as % of non-DM (blue) and DM (red) platelets in response to platelet agonists collagen 2 ug/ml, thrombin (IIa) 0.025 U/ml, U46619 10 uM, ADP 5 uM, and AA 0.5mg/mL. Aggregation to ADP and AA shown for patients with and without DM, with or without P2Y12 inhibitor (P2Y12i) or aspirin use, respectively. Mann-Whitney test. **B**. Secretion of ADAM Like Decysin 1 (ADAMDEC1) and Decorin (DCN) into the releasate of non-DM (blue) and DM (red) platelets following stimulation with low dose thrombin. Limma moderated t-test **C.** Glycated peptides identified by mass spectrometry of plasma from patients without or with diabetes. AA=arachidonic acid, U44619=thromboxane A2 receptor agonist.

**Supplementary Fig 2.**
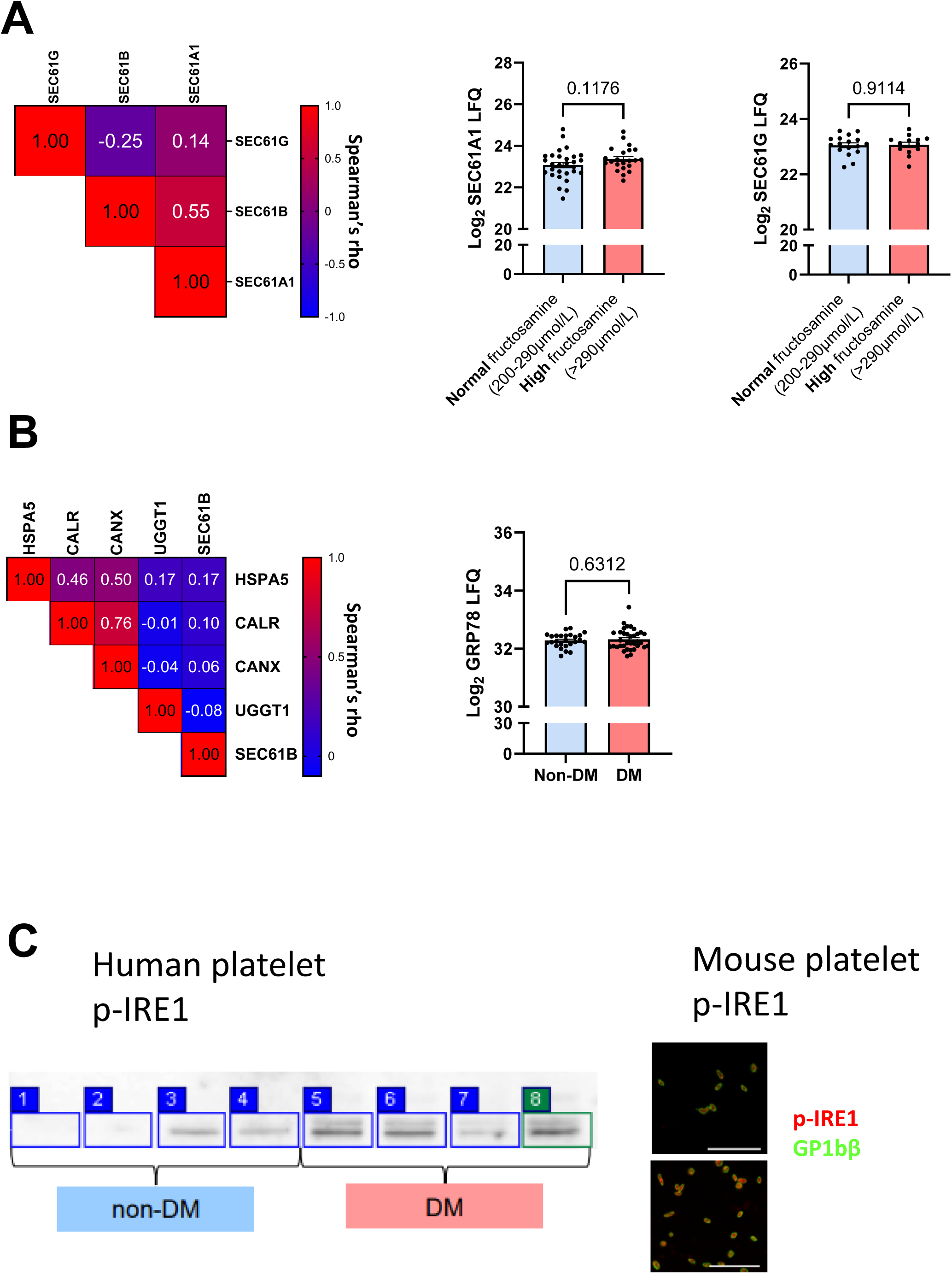
Increased abundance of SEC61B in diabetic platelets is not accompanied by increase in the abundance of the A1 and G subunits. **A.** Spearman’s rho correlation of the iBAQ abundance of the B subunit with the A1 and G subunits of the SEC translocon. Log2 LFQ of SEC61A1 and SEC61G in platelet lysate from patients with normal fructosamine (200-290 uM) versus high fructosamine (>290 uM); unpaired t-test with Welch’s correction **B.** Spearman’s rho correlation of SEC61B abundance with ER proteins HSPA5 (GRP78), CALR, CANX, UGGT1; Platelet lysate Log2 GRP78 LFQ by DM status **C.** Detection of p-IRE1 in resting lysates from non-DM and DM patients by western blot. Representative images. Detection of p-IRE1 in resting platelets isolated from non-hyperglycemic (Veh) and hyperglycemic (STZ) mice by immunofluorescent staining. Platelet receptor GPIbβ is shown in green and p-IRE1 in red. Representative images. Scale bar 20µm.

**Supplementary Fig 3.**
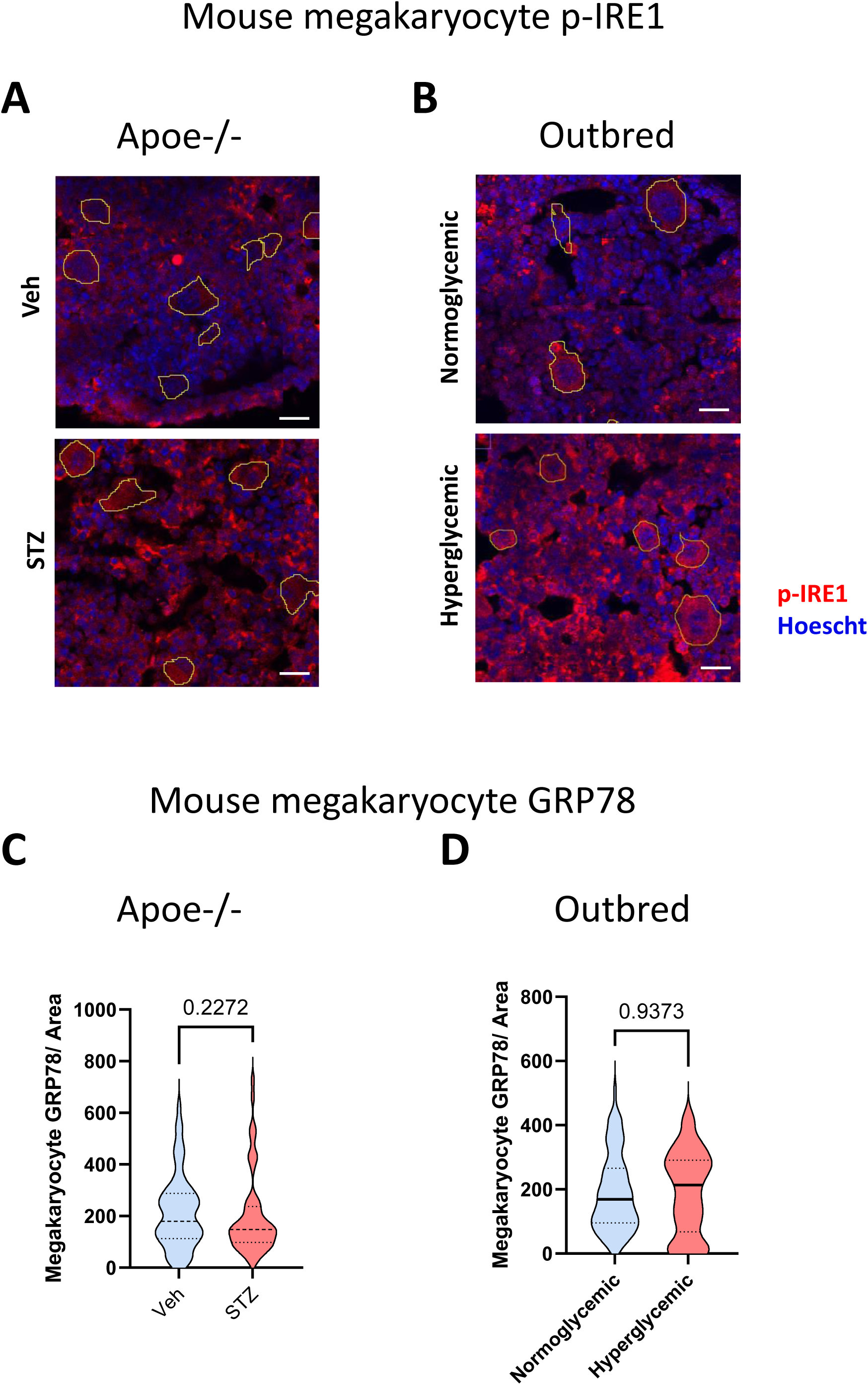
Increased phosphorylation of the IRE1 pathway in bone marrow megakaryocytes of diabetic mice is not accompanied by an increase in ER protein GRP78. **A.** Detection of p-IRE1 in megakaryocytes (outlined) of Apoe-/- with (STZ) or without (Veh) hyperglycemia and **B.** outbred mice with or without hyperglycemia by immunofluorescent staining. p-IRE1 is shown in red and nuclei in blue. Representative images. scale bar 20μm **C.** Immunofluorescence intensity of GRP78 in Apoe-/- mice without hyperglycemia (Veh, blue) versus with hyperglycemia (STZ, red) and **D.** outbred mice without hyperglycemia (blue) versus with hyperglycemia (red); Mann-Whitney test.

**Supplementary Figure 4.**
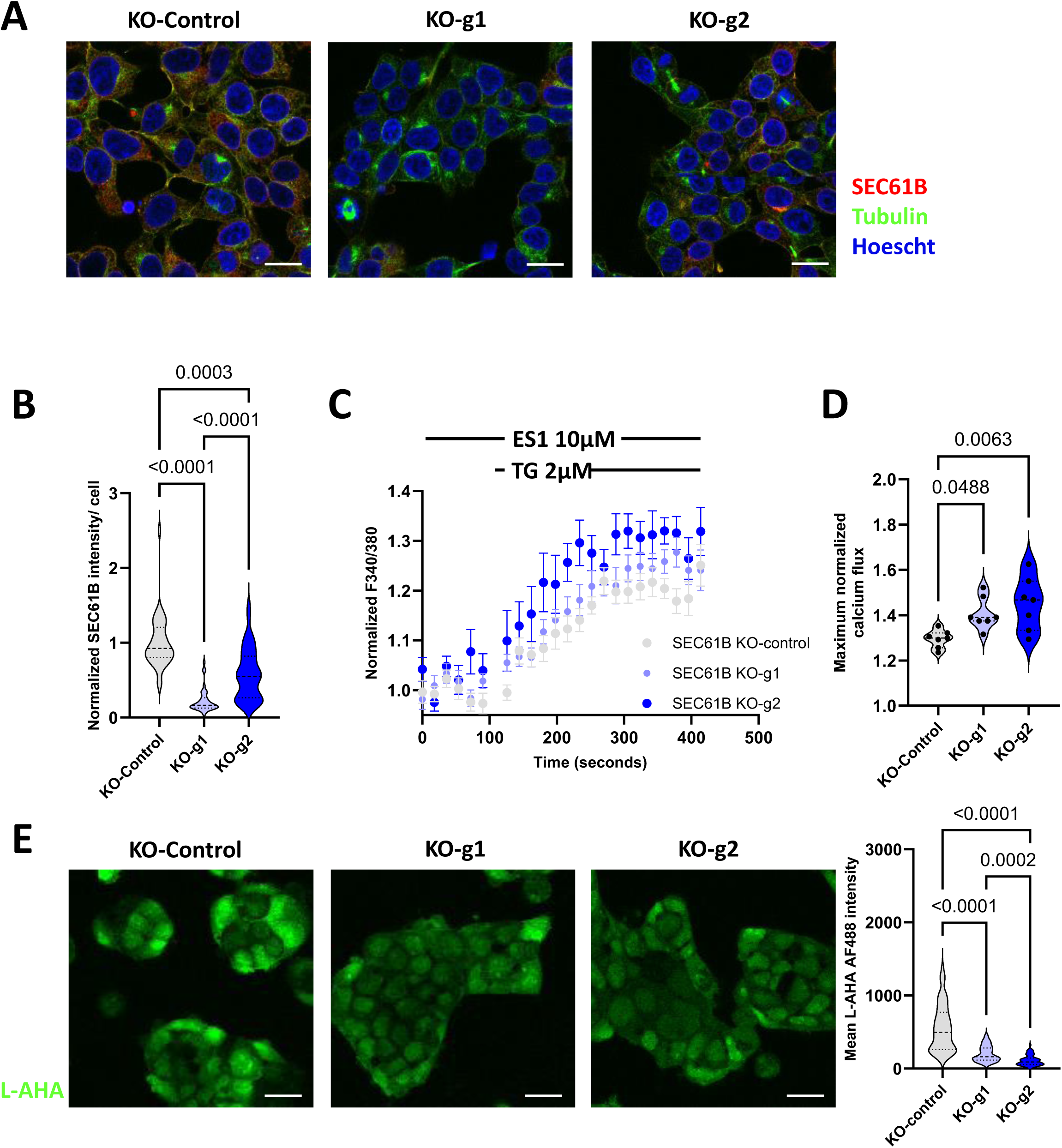
SEC61B depletion leads to increased ER calcium leak and reduced cellular protein synthesis. **A.** Representative images of SEC61B expression (red) in control and CRISPR/Cas9-mediated SEC61B knockout (KO), using two guide RNAs (g1 and g2), HEK293 cells. Tubulin in green and nuclear staining with Hoescht in blue. Scale bar 20µm **B.** Normalised SEC61B per cell; Kruskal-Wallis test **C.** Time course and **D.** peak normalised cytosolic calcium concentrations in control and KO cells (n=11 independent experiments, Ordinary one-way ANOVA). Cells were treated with eeyarestatin I (ES1) and thapsigargin (TG) to elicit the SEC61-mediated ER calcium leak. **E.** Representative images of protein synthesis (green) in control HEK293 cells, scale bar 20µm. **F.** Quantification of protein synthesis in the control and knockout cells (n=3 independent experiments), Kruskal-Wallis test.

**Supplementary Figure 5.**
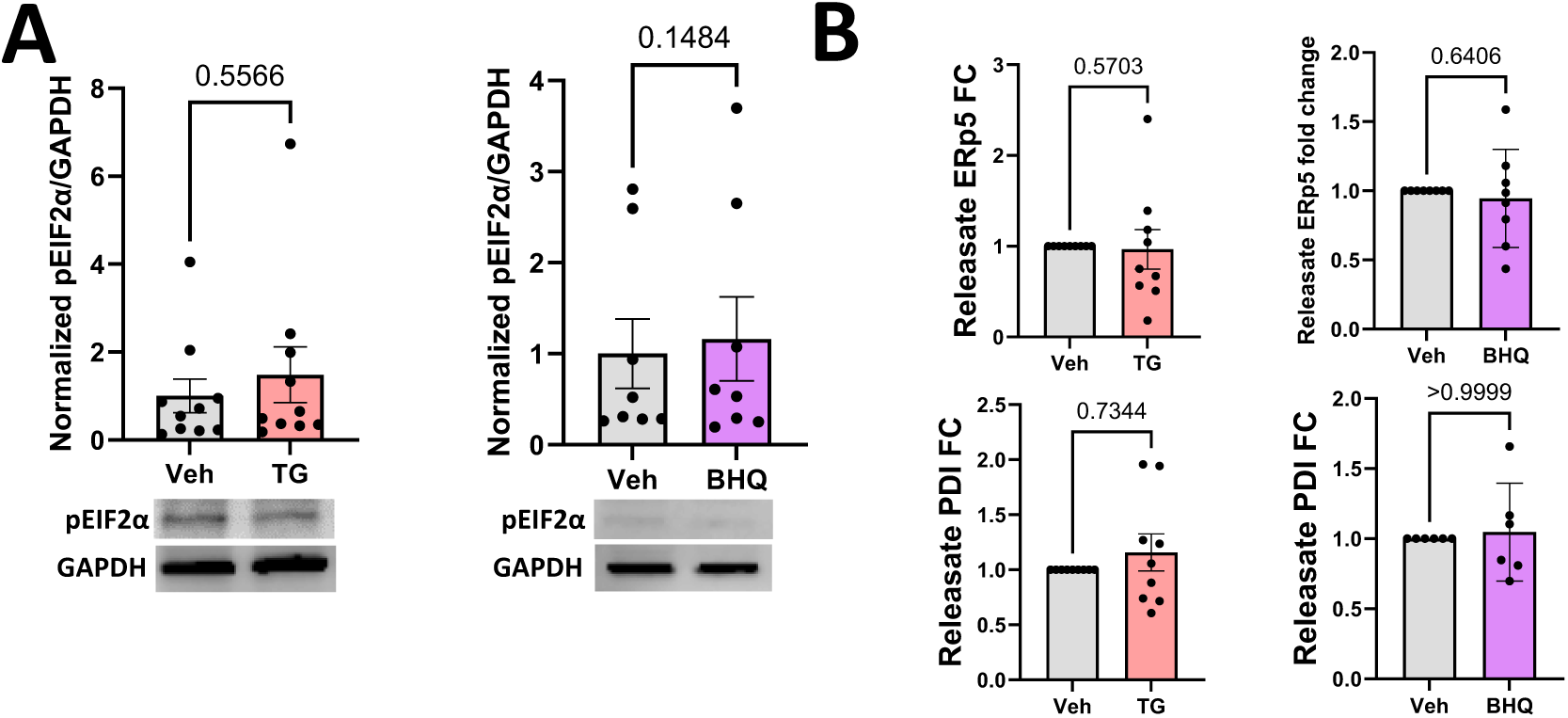
Stimulation of calcium efflux in human platelets by sarcoendoplasmic reticulum calcium ATPase inhibitors does not activate the PERK pathway and does not increase the secretion of PDI or ERp5 into the extracellular space. **A.** Ratio of phosphorylated eIF2a (p-eIF2a) to GAPDH of platelets treated with Veh (grey), TG (red) or BHQ (purple) **B.** Fold change of PDI and ERp5 in the platelet releasate after treatment with Veh (grey),TG (red), or BHQ (purple), as detected with Western blot. Wilcoxon test. BHQ: 2,5-di-t-butyl-1,4-benzohydroquinone; eIF2a: eukaryotic translation initiation factor 2A; PDI: protein disulfide isomerase; PERK: Protein kinase RNA-Like endoplasmic reticulum kinase; TG: thapsigargin.

